# Tissue-scale dynamic mapping of Hematopoietic Stem Cells and supportive niche cells in the fetal liver

**DOI:** 10.1101/2023.09.12.554625

**Authors:** Patrick M. Helbling, Anjali Vijaykumar, Alvaro Gomariz, Thomas Zerkatje, Karolina A. Zielińska, Kathrin Loosli, Stephan Isringhausen, Takashi Nagasawa, Ingo Roeder, Markus G. Manz, Tomomasa Yokomizo, César Nombela-Arrieta

**Author notes:** These authors contributed equally to the work (Patrick M. Helbling, Anjali Vijaykumar). Lead contact and correspondence: César Nombela-Arrieta, Department of Medical Oncology and Hematology University and University Hospital Zürich Häldeliweg 4, 8032, Zürich.

## Abstract

The fetal liver (FL) plays a fundamental role in the ontogeny of the hematopoietic system, by transiently providing a fertile microenvironment for the maturation, proliferation and expansion of fetal hematopoietic progenitors, as well as definitive hematopoietic stem cells (HSCs). Nonetheless, the cellular make up and identity of HSC niches in the FL remain poorly described. Here we employed a combination of 3D quantitative microscopy, bulk mRNA-seq, flow cytometry and cell-specific reporter models, to dissect the spatiotemporal dynamics of putative niche cells and HSCs in the FL microenvironment. We find that at peak stages of FL hematopoiesis, pro-hematopoietic cytokines are promiscuously expressed by endothelial, mesenchymal cells and most prominently by hepatoblasts. These multicellular consortia of parenchymal/stromal cells are spatially predominant in the FL, thus providing unrestricted access to supportive factors throughout the entire tissue. Accordingly, in these early phases HSCs are found scattered across the parenchyma not displaying obvious spatial biases within the broad microanatomy of the FL, but exhibiting clustering behaviors. This highly conducive microenvironment is transient and gets rapidly remodeled through hepatoblast differentiation and sterile inflammatory signaling, leading to the downregulation of hematopoietic factors and the contraction of supportive niches, which temporarily coincide with the exit of HSCs towards emergent BM tissues.

## Introduction

During embryonic development fetal liver (FL) tissues play major fundamental roles in the development of the hematopoietic system ^1^. In mice, the FL primordium is first seeded by an heterogeneous set of hematopoietic progenitors with limited differentiation potential generated in the yolk sac (FS), between days 7 and 9 of gestation (E7 and E9). YS-derived progenitors establish in the FL, where they differentiate and support hematopoietic function throughout fetal life ^2,3^. Shortly after (E11.5), the FL is additionally colonized by circulating multipotent progenitor cells, which previously emerge through endothelial to hematopoietic transition from the walls of large intra embryonic (IE) arterial vessels ^4–6^. Recent studies using sophisticated lineage-tracing mouse models demonstrate that within the FL microenvironment, progenitors originating in IE locations mature and simultaneously seed all layers of the definitive hematopoietic hierarchy, including long-term repopulating HSCs ^7^. While evidence suggests that most FL HSCs are poised to differentiation, the pool of transplantable HSCs with repopulating potential undergoes very active proliferation and expansion from E12.5 to E14.5 ^8–10^. After these critical phases, HSCs gradually migrate to the developing BM and spleen, a transition that coincides with the transformation of the FL into an epithelial tissue ^1^. Thus, the FL microenvironment acts as a crucial and unique signaling hub transiently instructing the maturation, self-renewal, proliferation and differentiation of the overlapping waves of progenitors and HSCs, which sustain fetal and eventually lifelong hematopoiesis ^11^.

As in other hematopoietic organs, HSCs in the FL are thought to reside in specific microanatomical locations termed niches, where extrinsic soluble and contact-dependent extrinsic cues from neighboring cells control their core functional properties and determine their fate ^12–14^. Numerous studies have focused in the dissection of BM niches in adulthood, where the majority of HSCs are preserved in a quiescent state and only enter cell cycle in a sporadic manner ^15–18^. However, our knowledge on the composition and location of pro-proliferative HSC niches in the FL remains incomplete. Akin to BM, evidence suggests that key microenvironmental regulatory cues controlling hematopoiesis are derived from stromal components of the FL. First, hepatoblasts, which are bipotent tissue-specific progenitors that give rise to hepatocytes and cholangiocytes, have demonstrated potential to directly regulate HSCs and hematopoiesis in both mice and humans ^19^. Hepatoblasts support and expand FL HSCs in vitro ^20,21^, presumably through their expression of a number of relevant cytokines, including Kit ligand (Kitl, also known as Stem cell factor Scf), as well as Angiopoietin-like proteins (Angptl) 2 and 3, Thrombopoietin (Tpo) and Eythropoietin (Epo) ^22^. However, through developmentally-timed differentiation *in vivo*, hepatoblasts rapidly lose identity, and only transiently coexist with HPCs and HSCs in the same tissue context. The temporal and spatial dynamics of hepatoblast differentiation, and their precise contribution to maturation and expansion of HSCs throughout crucial stages of FL maturation *in vivo* are yet unclear. Second, HSCs have been reported to intimately interact with endothelium during FL colonization and reside in relatively close proximity to endothelial cells (ECs) lining the sinusoidal vasculature of the FL, which potentially provide HSCs and fetal progenitors with survival cues^23–25^

Finally, several lines of evidence further point to a role of cells of mesenchymal origin, in potentially supporting HSC expansion in FL. For instance, fibroblast-like cell lines derived from E14 FLs and exhibiting an epithelial-to-mesenchymal transition (EMT) phenotype, expand HSCs from BM and FL *in vitro* ^26^. Recent studies have attempted to dissect the composition and function of this compartment *in vivo*. Co-expression of Nestin and NG2 marks a subset of mesenchymal stromal cells (MSCs), which are spatially restricted to the adventitial layers of large portal vessels and also express crucial hematopoietic factors ^27^. Targeted depletion of this population of pericytes using the *Ng2*-Cre mouse model was associated with a significant reduction of HSC numbers and their proliferative activity at late stages of FL hematopoiesis ^27^. Nonetheless, hematopoietic cytokines such as Kitl and Cxcl12 are also produced by stellate cells and their precursors, which likely constitute the largest fraction of MSCs in FL ^28^. Strikingly, depletion of *Kitl* in *Pdgfr*-Cre^+^ cells, which comprise at least both stellate cells and NG2 pericytes did not cause substantial effects on HSC and hematopoietic progenitor (HPC) subsets29.

Therefore, despite major progress in our understanding of the extrinsic regulation of hematopoiesis in hepatic tissues, a dynamic, comprehensive and spatial overview of the emergence, establishment remodeling and extinction of supportive HSC niches is currently lacking. While imaging studies have been instrumental in revealing the localization of HSCs and uncovering functional associations to partner cell types, the nuanced global quantitative mapping of HSCs has not been achieved in the FL ^30^. Progress in this direction has been largely hampered by the challenges associated to the reliable and specific identification of rare HSCs *in situ* ^31^.

Here we employ two reporter models to screen for and visualize putative major cellular players in HSC niches in the FL. We report that expression of *Cxcl12* and *Kitl* and most of the extrinsic factors thus far described to expand HSCs, is distributed across three major stromal/parenchymal elements. Using 3D imaging we analyze the spatiotemporal dynamics of these multicellular structures and describe the active remodeling of their transcriptomic signatures throughout the time window in which most active HSC differentiation and expansion has been reported to take place in the FL. Finally, we deliver the first tissue-wide global map of HSC and HPC distribution in FL tissues.

## Results

### Heterogeneous and promiscuous expression of supportive HSC niche factors in FL stromal cells

Expression of *Cxcl12* and *Kitl* distinctively marks HSC-supportive niche cells in adult BM tissues ^32–35^. Both cytokines have been reported to critically regulate HSC migration and expansion during embryonic stages and specifically in FL hematopoiesis ^35–37^. We therefore employed transgenic reporter *Cxcl12^GFP^* and *Kitl^GFP^* mouse models to screen for expression of these cytokines in stromal and hematopoietic elements of FL at E13.5, and thus mark putative niche cells at a developmental stage during which highly active HSC expansion takes place ^38^. Similar to BM tissues, expression of GFP was exclusively confined to the non-hematopoietic fraction of the FL (CD45^-^ Ter119^-^) in both reporter lines, as assessed by flow cytometry (FC) (Figure 1A and 1B). Highest expression levels of *Cxcl12^-^*GFP and *Kitl*-GFP were detected in non-endothelial CD45^-^Ter119^-^CD31^-^ stromal cells. Phenotypic characterization revealed that this fraction was heterogeneous and could be subdivided in at least two populations, which included i) a subset of cells of hepatic origin mostly consisting of immature DLK-1^+^CD249^+^CD140b^-^ hepatoblasts (HEP in figure panels), and ii) DLK-1^+^ CD140b^+^ cells, identified as mesenchymal stromal cells (MSCs) based on co-expression of CD51, CD166, and CD105 (Figure 1C and D). Albeit at lower levels, in both reporter mouse models GFP expression was also detected in hepatic endothelial cells (ECs), which co-expressed CD31, Lyve1, CD105, and CD144 (Figure 1C). At this stage of embryonic development all three subsets were relatively rare in cell suspensions from entire FL, collectively not exceeding 2% of the total cellular content (Figure 1E).

**Figure 1.**
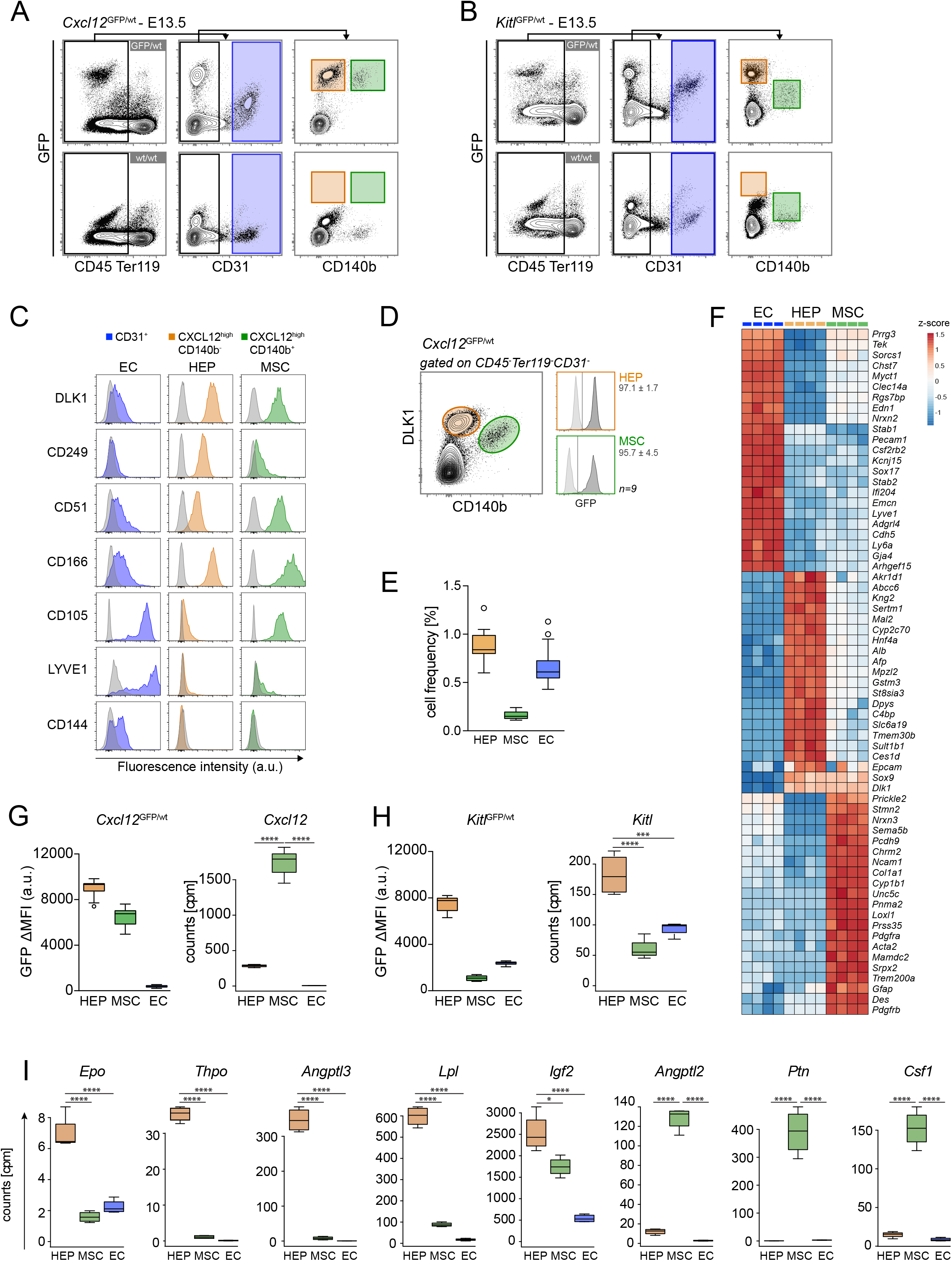
Heterogeneous and promiscuous expression of supportive HSC niche factors in FL stromal cells. (**A and B**)Flow cytometry gating strategy of E13.5 fetal liver stromal cells of *Cxcl12*^GFP/wt^ (**A**) and *Kitl*^GFP/wt^ (**B**) transgenic mice. GFP expression is restricted within the stromal compartment. Low GFP intensity is detected in CD45^-^Ter119^-^CD31^+^ endothelial cells (EC, blue gate), compared to GFP^hi^ in CD45^-^Ter119^-^CD31^-^CD140b^-^ (orange gate) and CD45^-^Ter119^-^CD31^-^CD140b^+^ mesenchymal stromal cells (MSC, green gate) cells. (**C**) Detailed immunophenotyping for the three defined stromal gates allow identification of CD45^-^Ter119^-^CD31^-^CD140b^-^ as hepatoblasts (HEP). (**D**) Histograms depict expression levels of *Cxcl12*-GFP on the gated HEP and MSC populations. Wild-type controls are shown in grey (representative data out of a total of n=9 embryos). (**E**) Quantification of the cellular frequency of the three stromal populations measured by flow cytometry in wild type B6 mice (n=13). (**F**) Heatmap depicting row normalized gene expression values of top 20 DEGs, as well as subset-specific genes of ECs, HEP and MSC cells confirming their identity. Four replicates are shown per subset (**G**) Quantified mean fluorescence intensity (MFI) of *Cxcl12*-GFP (n=13) and mRNA expression levels (n=4) of *Cxcl12* in all three defined stromal populations. (**H**) MFI of *Kitl*-GFP (n=9) and RNA expression levels (n=4) of *Kitl* for the defined stromal populations. (**I**) Expression levels of multiple genes linked to hematopoietic and HSC regulation in stromal populations by bulk mRNAseq. Data are shown as box plots depicting mean and 25 and 75 percentiles of 3 replicates for HEP and 4 replicates for MSC and EC. Statistical significance was analyzed using one-way ANOVA. *, P < 0.05; ***, P < 0.001; ****, P < 0.0001.

We employed the immunophenotypic signatures described above to purify ECs, MSCs and, hepatoblasts from E13.5 FL and perform bulk mRNA sequencing (Figure 1D and 1F). Differential expression analysis between all subsets broadly confirmed the identity of all populations. Hepatoblasts were characterized by highest expression of classical hepatic cell markers such as *Afp*, *Alb* or the transcription factor *Hnf4a*. In turn, MSCs were defined by expression of a large number of matrisomic genes, markers for hepatic stellate cell progenitors (*Des*, *Ncam1),* as well as *Pdgfra* and *Pdgfrb,* which had been employed as the cell surface marker for their isolation (Figure 1F). Finally, ECs expressed high transcript levels of pan-endothelial markers including *Pecam1*,*Tek*, *Cdh5* and *Enmc*, as well as sinusoidal (*Lyve-1*, *Stab1*, *Stab2)* or prototypical markers of arterial specification (*Gja4*, *Sox17,* and *Ly6a*), thus reflecting that the phenotypic signature employed for isolation did not discriminate between the different types of EC subtypes found in the developing FL ^39^. Both Gene Ontology (GO) terms and differentially expressed genes (DEGs) aligned well with the core functions described for all subsets. While hepatoblast-specific genes were very strongly enriched in several categories related to metabolic processes, MSC-specific signatures were overrepresented in genes linked to ECM organization connective tissue and mesenchymal development and axon guidance. As expected, EC-specific profiles were dominated by genes involved in wound healing, coagulation angiogenesis and immune regulation (Supplementary Figure 1). Collectively, these results validated the specificity of our isolation procedures and the identification of three major stromal components in early FL development. Of note, expression at the mRNA level of both *Cxcl12* and *Kitl* was confirmed in all three subsets, although transcript levels did not precisely mirror the expression patterns of the GFP reporter genes in both strains. While fluorescence intensity was brightest in hepatoblasts for both mouse models, mRNA levels of *Cxcl12* were highest in MSCs and comparatively almost negligible in ECs (Figure 1A, 1G). Nonetheless, for *Kitl* we observed a closer correspondence between mRNA transcript abundance and *Kitl*-GFP expression in the *Kitl*^GFP^ mice. In both cases, highest expression levels were found in hepatoblasts, but with a detectable and consistent expression in both MSCs and ECs (Figure 1B and 1H). We next mined our transcriptomic datasets for genes encoding factors previously involved in the regulation of HSCs, as well as in early lineage development and differentiation. Notably, at this developmental timepoint expression of several key factors was predominantly detected in HEP. Genes encoding for HSC-regulating factors such as *Igf2* and *Angptl3*, or *Lpl* were highly expressed in these cells, and detected at comparatively low levels in the other cell two types (Figure 1I). Notably, genes involved in erythro– and thrombopoietic differentiation (*Epo* and *Tpo)* could also be detected most prominently in hepatoblasts. Nonetheless, we also found that MSCs abundantly expressed other factors relevant to HSC and progenitor cell regulation. For instance, the expression of *Angptl2, Ptn* and *Mdk,* reported to influence HSC biology ^40,41^, as well as the myeloid cytokines *Csf1* and *Il34* was largely confined to this subset (Figure 1I and not shown). Altogether, these results suggested that the FL stromal microenvironment is formed by a complex multicellular consortium, which collectively provides a plethora of hematopoietic cytokines that fundamentally orchestrate FL hematopoiesis.

### Widespread spatial distribution of putative HSC niche cells within FL parenchyma

Flow-based quantifications suggested that all putative niche cells are relatively scarce in the FL (Figure 1E). Nonetheless, previous studies have shown that stromal subsets are notoriously difficult to extract into cell suspensions and are thus underrepresented in cytometric measurements ^42^. Therefore we next employed 3D imaging to visualize the abundance and spatial distribution of *Cxcl12*-GFP and *Kitl*-GFP expressing cells. For this purpose, we adapted a tissue processing and microscopy workflow, which we had previously optimized for BM imaging and that permits the generation of 3D images of thick, optically cleared tissue slices (∼200 µm) covering single entire FL lobes with single cell resolution (Figure 2A and Supplementary Movie 1) ^43^. Immunostaining allowed for tissue-scale visualization of vascular hepatic subtypes, including the sinusoidal tree and large portal vessels, marked by perivascular expression of smooth muscle actin (SMA). These analyses revealed a striking and conspicuous abundance of cells displaying variable expression of the *Cxcl12*-GFP transgene, which pervaded all liver regions, including perisinusoidal, periportal, and cortical areas (Figure 2B, 2C and Supplementary Movie 1). In line with flow cytometry analyses, various GFP^+^ cell types could be discriminated via marker expression and morphological criteria. CD140b^+^ mesenchymal MSCs displayed fibroblastic morphology and preferentially lied adjacent to sinusoidal vasculature (Figure 2C and 2D). In turn, CD140b^-^ cells expressing GFP at high intensities, exhibiting a nuclear cuboidal morphology and displaying dimmer staining for DNA, corresponded to hepatoblasts (Figure 2D), which were found globally scattered and densely populated the entire tissue parenchyma. Finally, EC liver sinusoidal vessels marked with Lyve-1 (not shown) expressed low to negligible levels of GFP. At this stage, vessel density and occupancy of FL tissues was very high, with sinusoids dominating the compartment and branching to all regions of the liver (Figure 2B).

**Figure 2.**
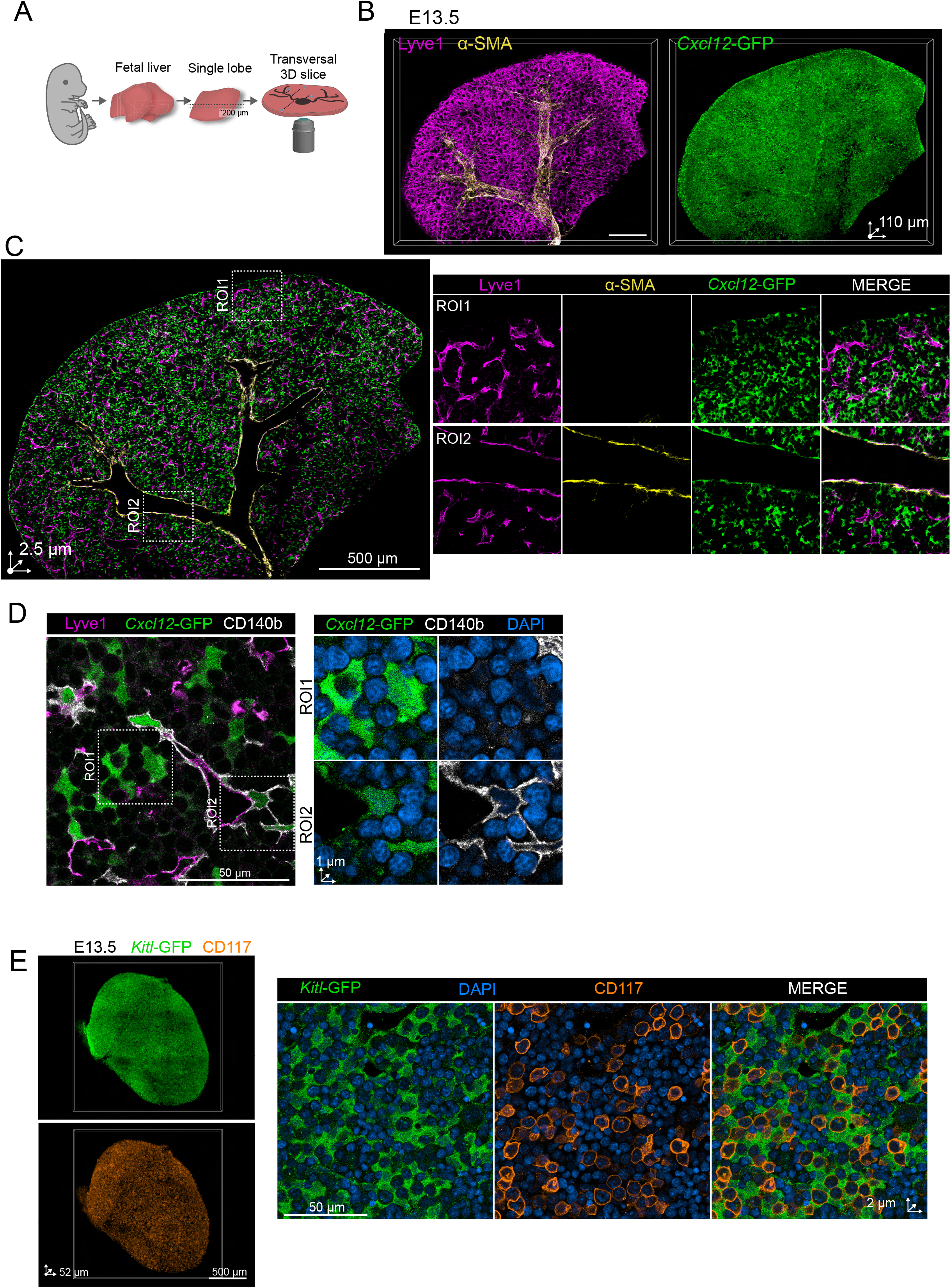
Spatial distribution of putative HSC niche cells within FL parenchyma. (**A**) Scheme depicting the three-dimensional quantitative microscopy (3D-QM) pipeline for FL imaging. FLs are dissected into individual lobes and sectioned into thick (250µm) slices, which are immunostained and optically cleared for confocal laser scanning microscopy. (**B**) 3D microscopy image of a slice covering a FL lobe (110 µm thickness) and (**C**) optical section of a whole slice (left) and regions of interest ROIs (right) of a E13.5 immunostained FL showing Cxcl12^GFP/wt^ expression (green) with sinusoidal vasculature labelled by Lyve1 (magenta) and arterial labelling by α-SMA (yellow). (**D**) High-resolution FL optical section showing CD140b-expressing (white) GFP^+^ (green) in Cxcl12^GFP/wt^ mice. CD140b-GFP-expressing populations with cuboidal-shaped and DAPI^dim^ nuclei correspond to HEP (**E**) 3D representation and high-resolution optical sections of an immunostained E13.5 fetal liver slice (52 µm thickness) showing global distribution of *Kitl*-GFP expression (green) and c-Kit^+^ hematopoietic progenitor cells (orange).

A comparable spatial and cellular expression pattern was observed in livers from *Kitl^GFP^* embryos at these stages (Figure 2F). Co-immunostaining of c-Kit^+^ HSPCs revealed a diffuse and widespread distribution of hematopoietic progenitors in FL. Notably, upon close inspection c-Kit^+^ HSPCs could be most often found in close association to, or fully embedded within large clusters of *Kitl-*GFP^+^ hepatoblatsts, suggesting direct contact-dependent functional interaction between both subsets (Figure 2E). Altogether, our observations suggest that, similar to what has been reported in BM tissues ^42^, the availability of HSPC-supporting factors is diffuse and spatially unrestricted in FL at early developmental timepoints, with almost all tissue resident cells localizing in close proximity to potentially critical niche components.

### Dynamic FL tissue remodeling results in the contraction of the pool of HSC-supporting niche components

HSC expansion takes place within a narrow developmental time window before hematopoiesis shifts towards BM and splenic tissues. Therefore, we investigated the quantitative and spatial dynamics of stromal cells in the FL parenchyma from early FL hematopoiesis to postnatal stages. From E13.5 until P1.5, the FL massively expands, as evidenced by substantial increases in organ weights (Figure 3A). Similarly, total FL cellularity drastically increases until E17.5, but total numbers of extracted cells drop at P1.5, which we explain by the exit of hematopoietic cells and the increased rigidity of tissues at this timepoint, which precludes full and efficient extraction of all stromal types (Figure 3B). Nonetheless, we detected a constant and gradual increase in the relative frequencies of ECs, which also translated into a substantial expansion of total EC numbers (Figure 3C and 3D). In turn, the overall frequencies and total cell numbers of MSCs remained low and largely constant during this developmental transition. Both ECs and MSCs maintained relatively stable levels of *Kitl*-GFP and *Cxcl12*-GFP until P1.5 (Figure 3E and 3F). Notably, cells falling under the phenotypic signature of hepatoblasts experienced a transient increase in numbers from E13.5 to E15.5, before undergoing a sharp decline until birth (Figure 3C and 3D). This virtual disappearance of hepatoblasts was explained by their differentiation into mature hepatocytes, which led to the downregulation of Dlk1 expression and their exclusion form the phenotypic gate used here to define the hepatoblast population (not shown). Most importantly, remaining hepatoblasts, and to a greater extent differentiated hepatocytes (Dlk-1^lo/-^) strongly downregulated *Kitl*-GFP expression (Figure 3E). Using qRT-PCR we confirmed that declining GFP expression in hepatoblasts was mirrored by progressively lower expression of *Kitl* transcripts in hepatoblasts from E13.5 to P1.5 (Supplementary Figure 2A). Most importantly, the collective downregulation of *Kitl* in the hepatocyte lineage translated into a major reduction in protein levels of KITL as measured by ELISA, thereby suggesting that hepatoblasts are the responsible stromal element producing this factor during this critical developmental phase (Supplementary Figure 2B). Of note, while acquisition of a mature phenotype led to decreased intensity of the *Cxcl12*-GFP, undifferentiated hepatoblasts maintained similar levels of expression at E17.5 compared to E13.5 (Figure 3F and Supplementary Figure 2A).

**Figure 3.**
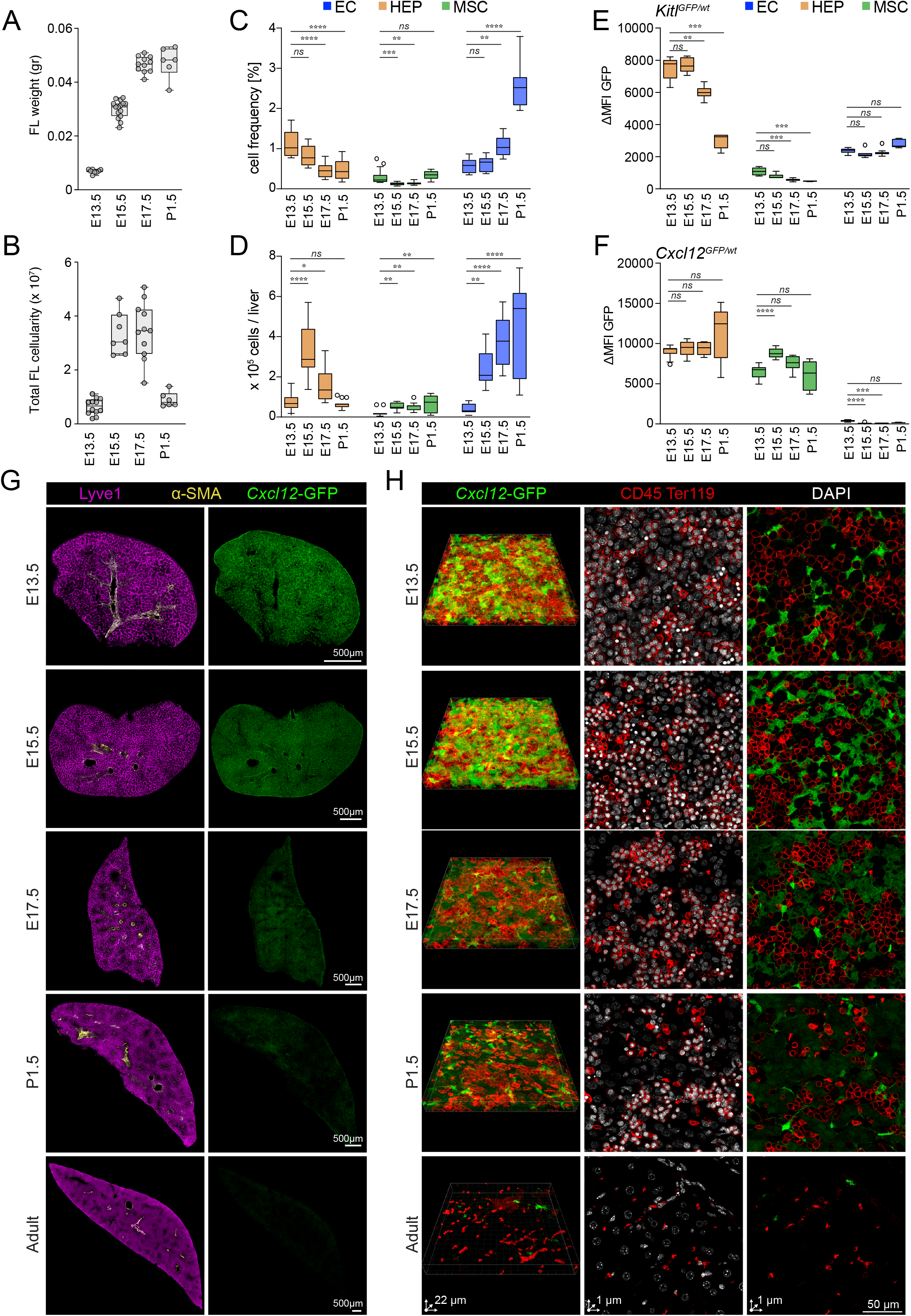
Dynamic FL tissue and HSC niche remodeling. FL weights (**A**) and total cellularity (**B**) at embryonic timepoints E13.5, E15.5, E17.5 and post-natal P1.5. (**C**) Dynamics of relative cell frequencies and absolute cell numbers (**D**) of HEP, MSCs and ECs subsets at different developmental timepoints (n= 15-16 embryos, from 5-7 dams). (**E**) variations in GFP expression as measured by flow cytometry over the same developmental timeframe in *Kitl*^GFP/+^ (n = 5-12 embryos, of 2-3 dams) and *Cxcl12*^GFP/+^ (**F**) (n = 6-14 embryos from 3-4 dams) mouse strains. Statistical significance was analyzed using the Kruskal-Wallis test with Dunn’s posttest. *, P < 0.05; **, P < 0.01; ***, P < 0.001, **** P < 0.0005. (**G**) 3D images of entire lobes representing developmental dynamics of organ-wide *Cxcl12*-GFP expression (green) over a developmental timeframe (E13.5-E17.5) and P1.5 and adult liver. Sinusoidal vasculature marked by expression of Lyve1 (magenta) and large vessels expressing α-SMA (yellow) are shown in left panels. (**H**) 3D images of restricted volumes (left) and 2D images of one single optical section (right panels) depicting variations in distribution and expression levels of *Cxcl12*-GFP (green) over the analyzed embryonic/postnatal timeframe, as well as in adult liver. The distribution of CD45^+^ and Ter119^+^ hematopoietic cells is shown in red. Representative images of at least three replicates are shown in **G** and **H**.

Histological examination using 3D imaging confirmed the marked global reductions of *Cxcl12*-GFP^hi^ cells, which appeared to gradually and homogeneously disappear throughout the entire FL parenchyma (Figure 3G, 3H and not shown). The contraction of the absolute surface covered by GFP signal was most pronounced between E15.5 and E17.5 and correlated with the spatial shrinkage of areas occupied by hematopoietic cell clusters (Figure 3H). In line with flow cytometry data, downregulation of GFP signals was mostly observed in cells with morphological characteristics of hepatoblasts, while MSCs remained as bright, scattered isolated cells. By early postnatal stages (P1.5) GFP^+^ cells were relatively rare, and progressively concentrated in periportal regions, and the subcapsular regions of the liver (Figure 3G, 3H and Supplementary Movie 2). As anticipated, similar temporal dynamics of GFP^+^ populations were observed in *Kitl*^wt/GFP^ mice (not shown). In summary, the stromal compartment of the FL undergoes a profound, developmentally-timed remodeling process that leads to the downscaling of niche cellular pools expressing crucial HSPC supporting factors, principally due to the differentiation of hepatoblasts into mature hepatocytes.

### Hematopoietic cytokine downregulation and inflammatory signaling underpin the remodeling of the FL microenvironment

We next sought to examine the global changes emerging in stromal cells at the transcriptomic level using bulk mRNA sequencing during the transition from a hematopoietic to a metabolic organ in the FL (Figure 4A). As shown in Figure 4B, principal component analysis (PCA) revealed substantial remodeling of transcriptomic profiles in hepatoblasts, MSCs and to a lower extent ECs from E13.5 to E17.5. We first focused on the changes in gene expression and pathways related to niche-dependent control of HSCs and progenitors. Transcript levels of *Kitl*, *Epo* and *Thpo,* which were most prominently expressed in hepatoblasts strongly decreased in this cell type; a trend that was confirmed using qRT-PCR (Figure 4C and Supplementary Figure 2B). Nonetheless, expression of other niche factors such as *Igf2*, *Angptl2*, *Anptl3*, *Ptn* and *Mdk* remained largely constant or even increased during this transition in the three different cell types (Figure 4C, Supplementary Figure 2B and data not shown). These data strongly suggest that within a short developmental period, transcriptomic remodeling of the hepatocyte lineage encompasses a relative decrease of HSC supportive potential, which could be at least partially compensated by expression of crucial factors by FL ECs and MSCs.

**Figure 4.**
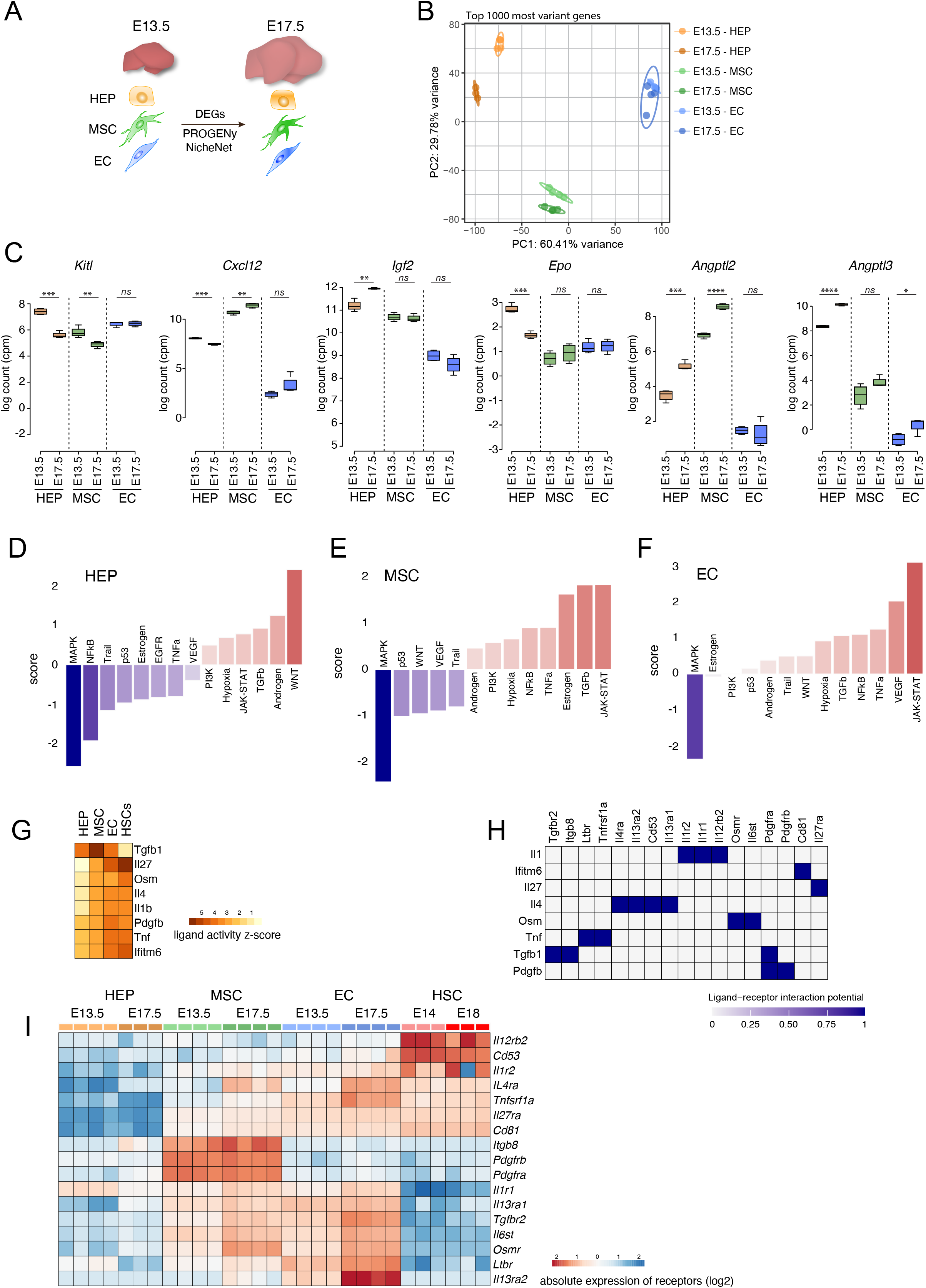
Inflammatory pathways drive intense transcriptomic remodeling of HEP, MSCs and ECs in FL during the early phases of mid-gestation. (**A**) Schematic representation of the developmental timepoints, cell types and computational models employed to analyze transcriptomic changes in FL between E13.5 and E17.5. (**B**) Principal component analysis (PCA) of HEP, MSC and EC cells isolated from FL at E13.5 and E17.5. (**C**) Changes in expression of crucial specific niche marker genes in HEP, MSC and EC. The plots show pooled data from three/four bulk RNA-seq experiments. Statistical significance was analyzed using unpaired Student’s *t* test. *, P < 0.05; **, P < 0.01; ***, P < 0.001, **** P < 0.0005. (**D-F**) PROGENy pathway analysis of differentially expressed genes (DEGs) in HEP, MSC and EC between E13.5 and E17.5. (**G-I**) NicheNet ligand-receptor analysis of DEGs in HEP, MSC, EC and HSC. (**G**) Potential upstream ligands driving differential expression between developmental timepoints in HEP, MSC, EC and HSC (from E14 and E18 datasets from ^46^). (**H**) Potential receptors expressed by HEP, MSC, EC and HSC associated with each potential ligand and (**I**) their expression levels.

To further investigate the potential effects of changes in gene expression on intracellular signaling we employed PROGENy, a computational tool, which allows to infer pathway activity in cells from RNA sequencing data based on consensus expression profiles derived from perturbation experiments ^44^. For each cell type sets of DEGs between E13.5 and E17.5 were computed (log_2_ fold change ≥ 1, FDR = 0.05) and activity scores were calculated for fourteen different pathways. In line with the reported roles of Wnt/b-catenin signaling in regulating hepatoblast specification towards cholangiocyte and hepatocyte lineages, the highest activity was detected in the WNT pathway in this cell type (Figure 4D). Of major interest, during this narrow transition, transcriptomic variations in most cell types were strongly related to the activation of inflammation-related signaling pathways. While TNFa and NFKB pathways appeared active in both MSC and ECs, the JAK-STAT pathway was detected in all stromal components among the highest activation scores (Figure 4D, 4E and 4F). Finally, changes in transcriptional programs revealed the common activation of the TGFb pathway in all three stromal components.

We next applied the NicheNet algorithm to gain insight into the ligands and receptors involved in driving these developmentally timed changes in gene expression ^45^. In accordance with the results generated by PROGENy, both Tgfb1 and TNF were ranked within the eight top potential ligands inducing transcriptomic remodeling of stromal cells (Figure 4G). Inflammatory ligands such as Il1, Ifitm6, Il4 and Il27, as well as, Pdgfb completed the list of top prioritized ligands. Notably, similar ligands were found to induce gene expression changes in FL HSCs isolated at E14 and E18 analyzing previously published RNA-seq data ^46^. Crucially, expression of TNF receptors such as Ltbr or Tnfsrf1a, or known receptors for IL1, IL4 and IL27, could be detected in HSCs, ECs and MSCs but not in hepatoblasts. Of interest, the tetraspanins CD53 and CD81, which have been implicated in the protection of HSCs from inflammatory stress, were expressed by FL HSCs and identified to respectively mediate the activity of Il4 and Ifitm6 on HSCs (Figure 4G, 4H and 4I) ^47,48^. Finally, Oncostatin M (Osmr), which is produced by FL macrophages and contributes to hepatocyte differentiation, was also ranked among the ligands with highest potential ^49,50^. Altogether, these results reinforce the notion that the downregulation of trophic HSC factors together with basal tonic inflammation acting on both the tissue microenvironment and HSCs, potentially jointly contribute to the tissue remodeling process that gradually drives cancelation of the niche and HSC exit from FL at these stages.

### Visualization and quantification of FL HSCs and HSPCs using a dual reporter mouse model

Given the scattered distribution of HSC supporting cells and their lack of spatial confinement we next set out to perform a comprehensive and global analysis of the localization of definitive HSCs and progenitors in FL and their dynamics throughout development. While complex immunophenotyping strategies have been used in the past to tracks HSCs *in situ*, high fidelity detection of rare HSCs has been hampered by the need to stain multiple markers in such approaches, which is especially challenging in the case of thick samples required for deep tissue imaging ^31^. Thus, we generated a novel dual reporter mouse model by crossing two previously published transgenic strains that allow for the endogenous labelling of HSCs and a broader hematopoietic progenitor (HPC) population. In the FL and BM of *Hlf^tdTom^* mice, expression of tdTomato (tdTom) specifically marks a fraction of c-kit^+^ progenitors and thus enables visualization of a heterogeneous subpopulation comprising various primitive HPCs (Figure 5A and Supplementary Figure 3B and Supplementary Movie 3) ^7,51^. To further distinguish HSCs among the targeted HPC subset, we crossed *Hlf*^tdTom^ mice to *Ctnnal1^GFP^* strain in which expression of GFP has been reported as highly specific to rare, non-dividing quiescent stem cells in the hematopoietic compartment of adult BM ^52^. In line with this reported HSC-specific labeling, we found that within FL, the expression of *Ctnnal*1^GFP^ was also largely confined to the HSC-enriched LSKCD48^-^CD150^+^ subset, with 88.3 ± 4.97 % of this phenotypically defined population carrying the GFP label at E13.5 in the FL (Figure 5A). Expression of GFP in HSCs also correlated with highest expression levels of Hlf^tdTom^ cells (Figure 5A and Supplementary Figure 3A). Most importantly, the majority of GFP^+^tdTom^+^ cells in this dual labelling approach were LSKCD48^-^ cells, and 51 ± 10 % (at E13.5) or 87 ± 3 % (at E17.5) were additionally CD150^+^ (Figure 5B) using very rigorous gating criteria. Of relevance, expression of *Ctnnal1*-GFP could also be detected, albeit at lower levels in the stromal compartment including a small fraction of cells of mesenchymal origin (CD45-Ter119-CD31-CD140b^+^, and a subset of CD31^+^ ECs (Supplementary Figure 3C and 3D). Imaging revealed that within the vascular compartment, *Ctnnal1*-GFP expression was restricted to ECs lining portal vessels (Supplementary Figure 3E). Therefore, the *Ctnnal*1^GFP^*Hlf*^tdTom^ double reporter mice allowed us to visualize and map large numbers of GFP^+^tdTom^+^ HSCs and GFP^-^tdTom^+^ HPCs, effectively discriminating them from other hematopoietic and non-hematopoietic GFP^+^ cells, as well as from spurious fluorescent signals with a very high level of confidence (Figure 5D).

**Figure 5.**
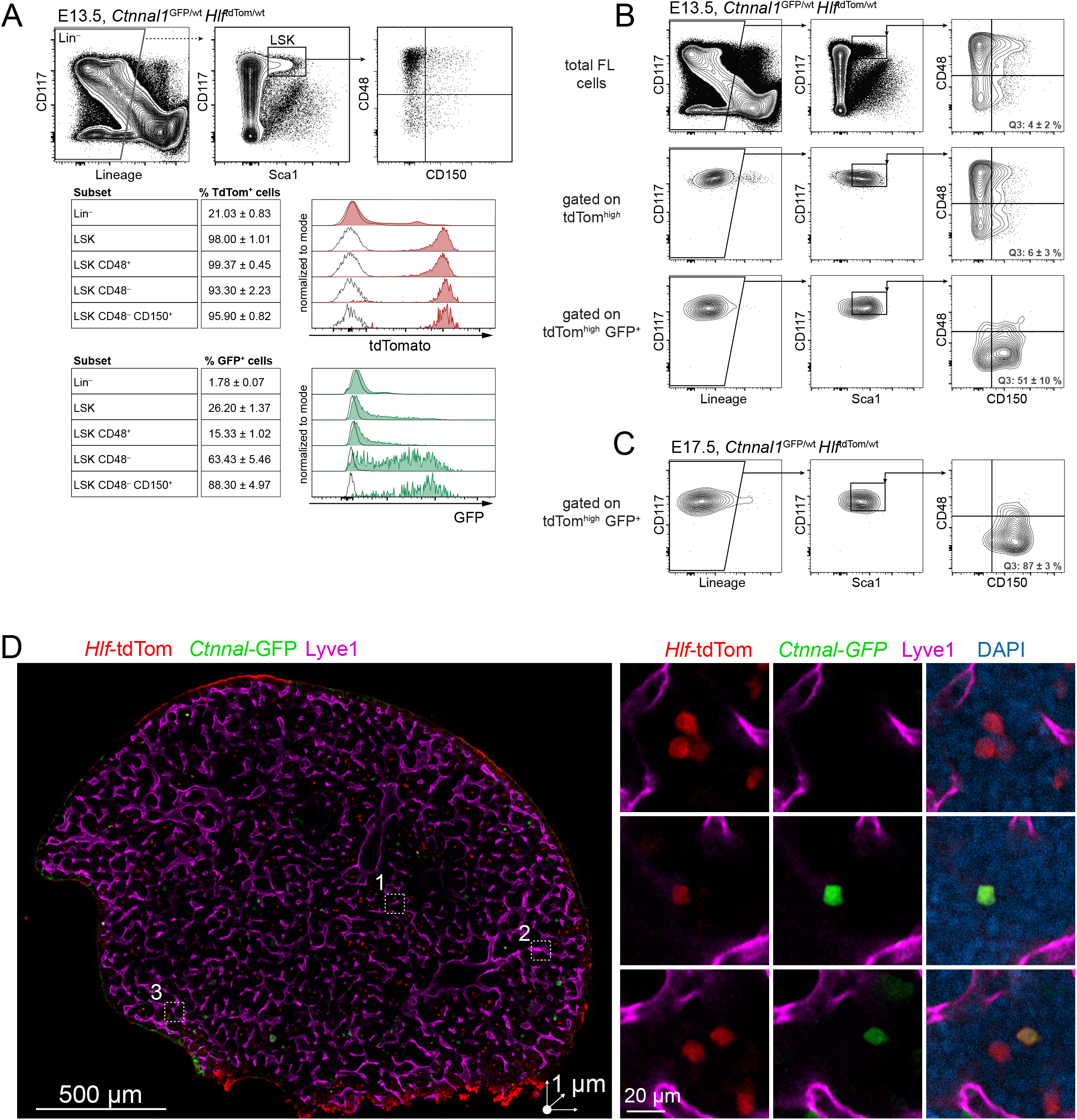
Specific labeling of HSCs and HSPCs for FC and 3D imaging using the *Ctnnal1*^GFP/wt^ *Hlf*^tdTom/wt^ dual reporter mouse model. (**A**) FC analysis of dissociated livers at E13.5 of *Ctnnal1*^GFP/wt^ *Hlf*^tdTom/wt^ double transgenic mice. Gating strategy of different HPC and HSC subsets to test for expression of both transgenes (top). Lower panels show the histograms of expression levels of both transgenes (right) and frequency of tdTom^+^ cells and GFP^+^ cells in each of the cellular fractions indicated. (**B and C**) Backgating of tdTom^high^ and tdTom^high^ GFP^+^ cells shows specific labeling of HPCs and HSCs, respectively in FL from E13.5 (**B**, n= 3) and E17.5 (**C**, n=5). Double positive cells tdTom^high^ GFP^+^ are highly enriched for the phenotypic stem cell signature LSKCD150^+^CD48^-^ (**D**) Optical section of an immunostained FL lobe from a *Ctnnal1*^GFP/wt^ *Hlf*^tdTom/wt^ embryo at E13.5. Vasculature is labelled by Lyve1 (magenta), tdTom in red and GFP in green. Right panels show regions of interest (ROI) depicting a cluster of HPCs (top panel, tdTom^+^GFP^-^) a single HSC (middle, tdTom^+^GFP^+^) and a HSC and HPC lying in close proximity (bottom).

### Widespread distribution of HSCs and HPCs throughout entire FL lobes

We next employed the labeling strategy described above and our previously described 3D quantitative imaging pipeline to perform a nuanced, tissue-scale analysis of the global distribution and interactions with blood vessels of HSCs and HPCs throughout FL parenchyma and across developmental stages. As described before, volumetric imaging datasets were generated by imaging thick tissue slices (50-125 µm thickness) of entire lobes. Slices were stained with the DNA dye DAPI to label all individual nuclei, and delimit tissue boundaries through the creation of a tissue mask (Supplementary Figure 4A and Supplementary Video 4). In turn, all vascular structures were stained with a combination of endothelial markers Lyve1 and Endomucin (Emcn). Segmentation of intravascular volumes in immunostained slices was automatically performed using a previously described deep learning computational model, which was trained with manually annotated ground truth data (Supplementary Figure 4B) ^53^. The segmentation metrics of this model for FL microvasculature has been previously reported ^53^. Subsequently, all segmented vascular structures were manually classified as *sinusoidal* or non-sinusoidal, termed here *large vessels*, which included portal and central veins readily identified based on size of luminal cross-section, on morphological features and on the presence of an ensheathing cell layer of alpha-smooth muscle actin-expressing cells (αSMA) (Figure 6A and 6B). At E13.5 intravascular spaces occupied up to 20.25 ± 3.24 % of the total hepatic volume, of which 14.18 ± 3.24 % corresponded to intrasinusoidal volumes, while the rest (6.06± 2.97 %) was occupied by large vessels. This pervasive vascularization pattern was maintained during development and organ expansion, as the volumetric fraction of vessels only experienced a subtle contraction by E17.5 (Figure 6C).

**Figure 6.**
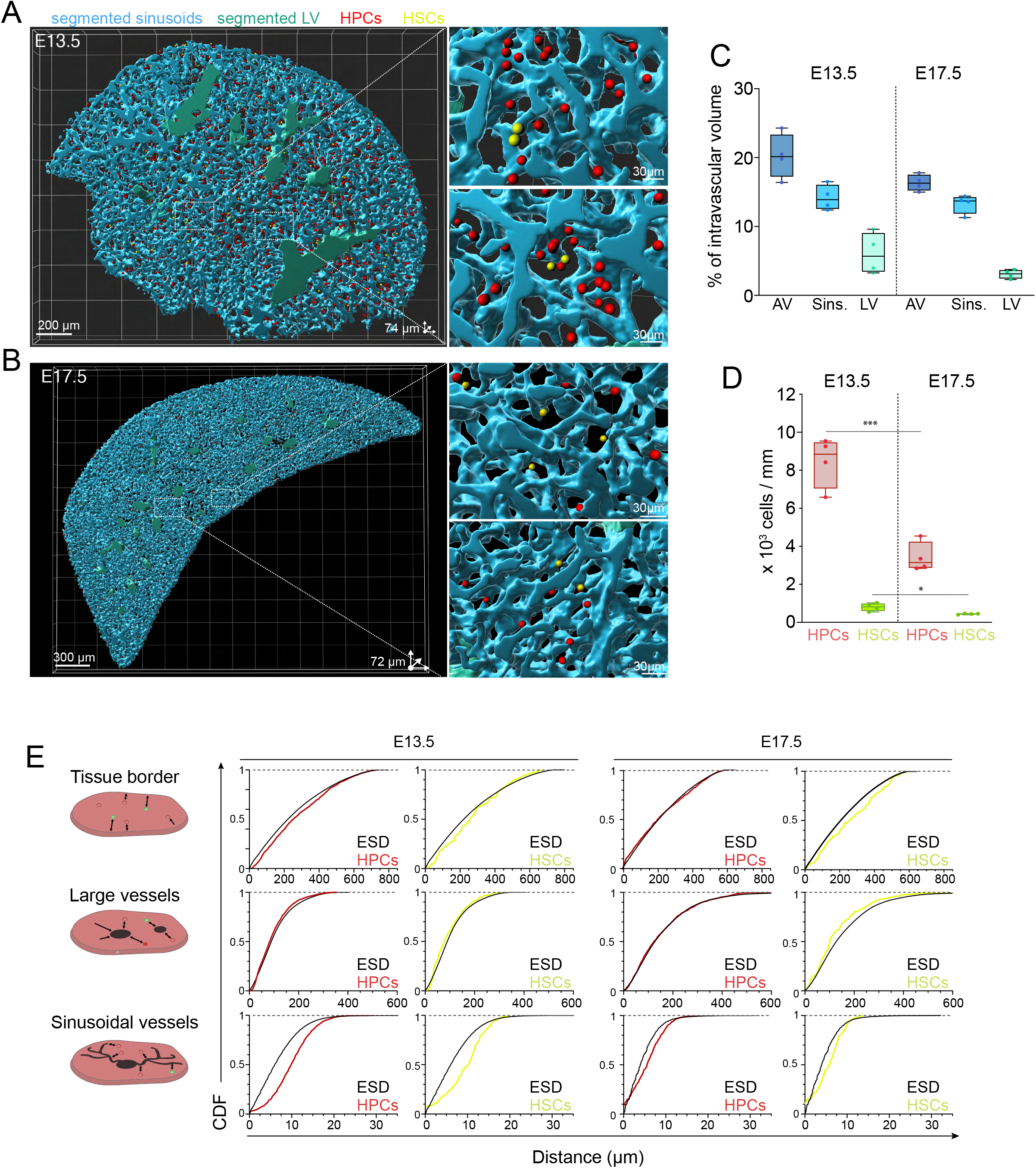
Hematopoietic stem cells distribute throughout the entire liver lobe and localize away from sinusoidal vasculature. (**A and B**) Representative computational reconstructions of slices of entire FL lobes and zoomed-in regions (right) from E13.5 (**A,** image generated from same dataset as in Figure 5D) and E17.5 (**B**) displaying segmented sinusoidal vessels (light cyan) and large vasculature (light green) and manually annotated and curated tdTom^+^GFP^-^ HPCs (marked by red spots) and tdTom^+^GFP^+^ HSCs (yellow). Size of scalebars and image depth is included in the figure panels. (**C**) Quantification of the fraction of intravascular volume out of the entire volume imaged corresponding to all vessels (AV), sinusoids (Sins) and large vessels (LV) at both developmental timepoints. Box plots depict maximum and minimum values, mean and all individual data points (n=4 embryos per developmental timepoint). No statistical significance was detected when comparing vessel volumes between timepoints with the Mann-Whitney *U* test. (**D**) Cellular density of HPCs and HSCs in FL 3D images from E13.5 and E17.5. Box plots depict maximum and minimum values, mean and all individual datapoints (n=4 embryos per developmental timepoint). Statistical significance was analyzed by unpaired Student’s *t* test *, P < 0.05; ***, P < 0.001.(**E**) Spatial analyses using points processes of the distribution of HPCs and HSCs with respect to the tissue border, to large vessels and to sinusoidal vessels at both E13.5 and E17.5. The 3D ESD (black lines) represents the distribution of distances from every voxel in extravascular spaces to the closest tissue boundary, large vessel or sinusoidal vessel in the tissue volume analyzed. The ESD is compared to the cumulative distribution function of the distances of every HPC (red) or HSC (yellow) in the tissue volume analyzed to the same anatomical landmarks. Graphs depict analysis of one individual representative tissue slice per timepoint out of a total of 4 analyzed per timepoint and cell type.

We manually annotated a total of 12332 HPCs (tdTom^+^ GFP^-^) and 1283 HSCs (tdTom^+^ GFP^+^) across a total of 8 FL slices (50-150 µm thick, median thickness 75 µm), four for each developmental timepoint (E13.5 and E17.5) investigated. Cells of interest were annotated based on cell size, round shape and intensity profiles. For quality control purposes, we quantified the fluorescence intensity in the GFP, tdTom and DAPI channels for a total of 2180 HPCs and 179 HSCs, which were randomly selected (∼250 HPCs and 20 HSCs per slice). In line with flow cytometry based measurements, cells classified as HSCs not only displayed high intensity GFP levels but were also consistently brighter than HPCs in the tdTom channel, while intensity in the DAPI channel was comparable between both subsets (Supplementary 5A, 5B and 5C). As expected, the observed cellular density of both cell types was substantially higher at E13.5 size compared to E17.5 FLs (Figure 6D). The reduction in the relative abundance of HSPCs at E17.5 is likely explained by declining nature of extramedullary hematopoiesis towards late fetal stages and the substantial increase in the size of the liver parenchyma.

Using all computational representations, i.e. (i) tissue mask, (ii) segmented sinusoidal and large vasculature, and (iii) annotated progenitor and HSC populations, we examined the presence of non-stochastic distribution patterns and associations to vascular structures of HSCs and HPCs with the statistical framework points processes, as previously done in the BM ^42^. We automatically computed the cumulative distribution function (CDF) of the distances of each cell (HSC or HPC) to the nearest structure of interest. The CDFs for both subsets were then compared to the distribution of the empty space distance function (ESD), which is given by the distances of every single voxel in the extravascular space to the same structures. The ESD thus provides a reference of how anatomical landmarks constrain the available territory for cells to reside in a certain tissue microarchitecture, and defines the null distribution, that is the one that cells would adopt if stochastically localized. In line with the widespread localization observed for supporting cell types, HPCs and HSCs at E13.5 were found scattered throughout the FL parenchyma and did not exhibit pronounced patterns of spatial compartmentalization at a global tissue scale (Figure 6A and Supplementary Figure 5D). Spatial analyses with respect to tissue boundaries revealed only a subtle decrease in the presence of both subsets in most outer regions of the liver (≤ 400µm from tissue boundary) (Figure 6E). Contrary to previous reports ^54^, HSCs did not display a preferential association to large vessels. Most importantly, the fraction of both cell types in direct contact or very close proximity (10-15 µm) to sinusoidal surfaces was lower than would be expected from a random distribution, which suggests no specific affinity to perisinusoidal sites (Figure 6E). Indeed, visual inspection confirmed that most HSCs and HPCs did not lie in direct contact with endothelial linings (Figure 6A, 6E and Figure 5C). Nevertheless, due to the spatial predominance of sinusoids, all HSCs and HPCs localized within very short distances (≤ 20µm) of sinusoidal vessels. Despite the reduction in overall HPC and HSC density, the described localization patterns were largely maintained at later timepoints and similar trends were observed at E17.5 (Figure 6E). Altogether we provide strong evidence that HSCs and the broader population of HPCs do not exhibit major spatial biases in FL and are amply distributed throughout the entire tissue parenchyma.

### HSCs and HPCs spatially associate in clusters during active proliferative phases

Visual analyses of our annotated datasets indicated the frequent presence of small clusters formed by multiple HPCs and most conspicuously HSCs, which were not necessarily next to each other but concentrated within limited tissue volumes (Figure 7A). To statistically test for potential significant clustering patterns of both cell types we computed the distance of each HSC or HPC to its nearest counterpart (belonging to the same subset), also termed nearest neighbor distance (NN, Figure 7B). The distribution of NN distances was plotted for each tissue slice and compared to the range of NN distributions obtained from 500 random simulations (Figure 7C). Of interest, this analysis uncovered that at E13.5 both HSCs and HPCs reside closer to cells of their own subpopulation than would be expected from a stochastic distribution (Figure 7C and 7E). Despite the scarcity of HSCs, this trend was conserved in all tissue slices analyzed and was detected up to distances of 100 µm between HSCs (Figure 7C). As an example, we found that 20-25 % of HSCs localized within less than 25 µm of its NN, a fraction that was 4-5 fold higher than that obtained from all random simulations (Figure 7D). Albeit HPCs populated FL at much higher densities, a similar clustering trend was detected in this subset (Figure 7E and 7F). Of note, this spatial patterning was strongly attenuated or virtually absent for both cell types at E17.5 (Figure 7C, 7D, 7E and 7F). Altogether, these analysis uncover an unexpected trend of non-stochastic spatial clustering during the early developmental phases in which expansion of HSCs and fetal progenitor cells in the FL is most active.

**Figure 7.**
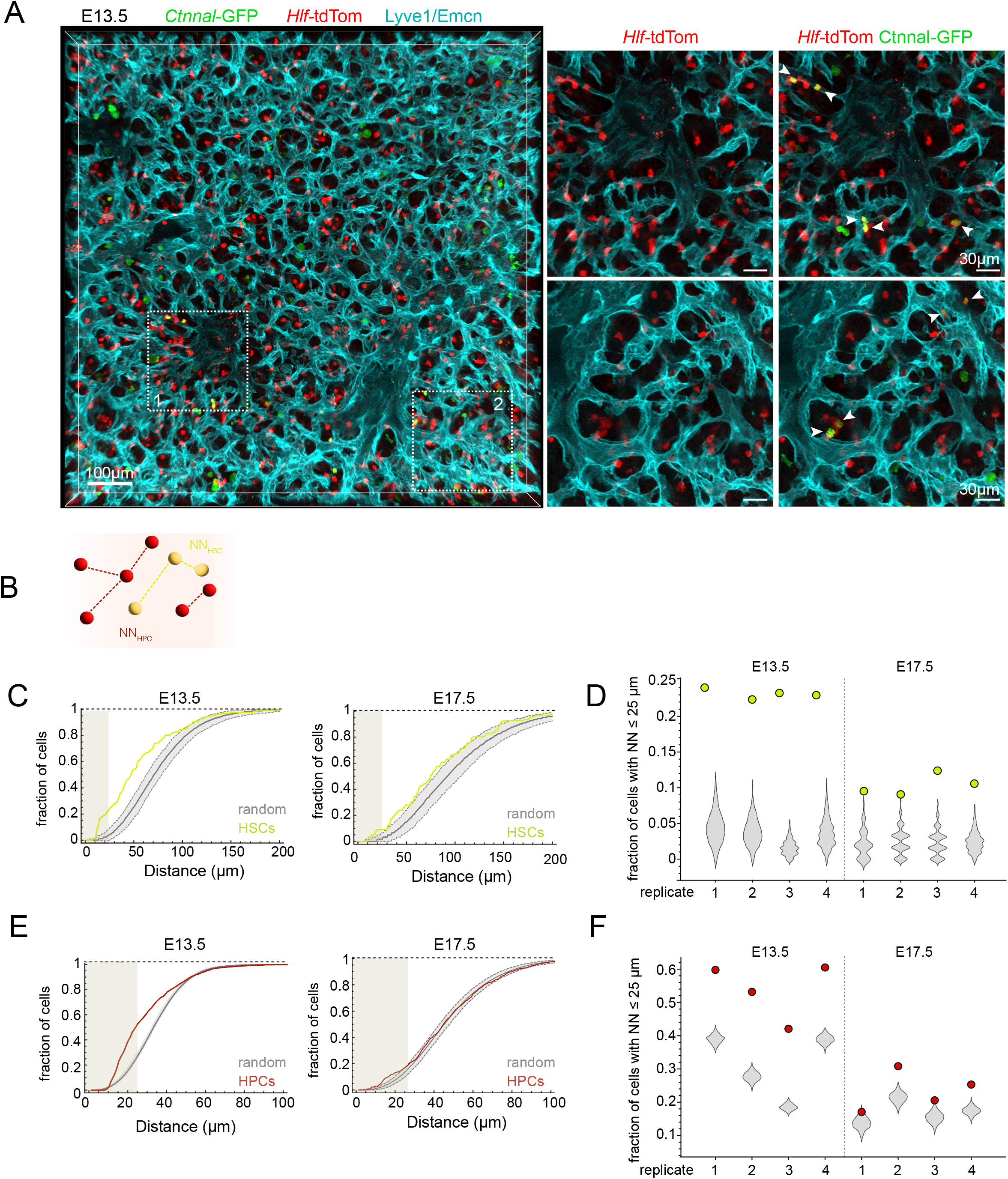
HSCs and HPCs spatially associate in clusters during active proliferative phases. (**A**) 3D image covering a large tissue region of a FL slice from a *Ctnnal1*^GFP/wt^ *Hlf*^tdTom/wt^ embryo at E13.5. Right panels show ROIs depicting areas of high density of tdTom^+^GFP^-^ HPCs and containing multiple tdTom^+^GFP^+^ HSCs. All blood vessels are marked in cyan labeled. (**B**) Schematic representation of nearest neighbor analysis. (**C, E**) Spatial analysis of distribution of nearest Neighbor (NN) distances of HSCs and HPCs compared to multiple random spot simulations in FL from E13.5 and E17.5 embryos. The CDFs of the NN distances of HSCs (**C**) and HPCs (**E**) are shown in yellow and red, respectively. Grey lines and envelopes represent the mean and standard deviation of 500 simulations performed within the same tissue volumes. Graphs depict analysis of one individual representative tissue slice per timepoint out of a total of 4 analyzed per timepoint and cell type. Shaded areas cover the distance of 25 µm to NN, shown in panels (**D, F**). Values of the fraction of HSCs (**D**) and HPCs (**F**) with one NN within ≤ 25 µm for 4 individual replicates per cell type and timepoint. Violin plots show the distribution of the values obtained by 500 random simulations using the same number of cells found in each tissue volume.

## Discussion

The detailed dissection of the identity and composition of HSC niches potentially holds the key for the informed design of novel strategies to efficiently maintain and expand HSCs ex vivo. ^55,56^. Nonetheless, while adult pro-quiescent HSC niches have been investigated in great depth, fewer studies have focused on understanding the extrinsic cues driving substantial and uninterrupted proliferation of HSCs in the FL microenvironment. In this study we combined cell-specific reporter mouse models with advanced 3D microscopy strategies and global gene expression analyses to visualize putative niche components and HSCs, map their spatiotemporal dynamics within the FL microarchitecture and analyze the structural and functional remodeling of the tissue microenvironment during the fetal time-window of highest hematopoietic activity.

We find that at E13.5 the expression of the most relevant cytokines reported so far to control FL hematopoiesis is heterogeneously distributed across three major subsets of the non-hematopoietic stromal compartment, which permeate the entire FL. Our results support the fundamental idea that during the peak expansion phases of HSCs, niche activity is not spatially confined, but accessible to virtually all FL residing cells irrespective of their specific microanatomical location. Indeed, MSCs and especially hepatoblasts and ECs are highly abundant and densely populate the FL, saturating all regions of the liver parenchyma. This is illustrated by the fact that hepatoblasts ubiquitously occupy the FL, and that all the extravascular space is contained within less than 20 µm of the nearest sinusoidal surface. These observations are in line with what we and others have described in BM tissues, where the abundance and spatial coverage of the key niche cellular networks is also exceedingly high and their proximity to HSCs is dictated merely by spatial constraints and not necessarily driven by attractive forces ^42,57^. Mirroring what we observe for niche cells, HSCs and HPCs exhibit no predilection towards residing in specific regions and follow a widespread and stochastic distribution with respect to major histological landmarks. Collectively, these observations strongly suggest that niche support is a general property disseminated throughout the whole FL, as opposed to previous studies in which analysis of reduced numbers of HSCs labeled within limited histological regions indicated a preferential residence HSCs in the proximity of periportal regions ^27^. Strikingly, using rigorous spatial statistical methods, we reveal that both HSCs and HPCs distribute according to clustering patterns, which may be explained either by the proximity between recently divided HSCs and HPCs or by the existence of local spatial affinities towards a yet undefined cellular subtype with highest niche activity. Therefore, further studies are needed to understand the functional implications of clustering during these stages.

Nonetheless, the described tissue topography, characterized by an unrestricted access to cells with HSC supportive properties, is transient and undergoes rapid remodeling between E13.5 and E17.5, which results in the drastic reduction of the pool of stromal supportive cells and the gradual contraction of the overall space they occupy in the FL. This is principally driven by the developmentally timed loss of immature hepatoblasts due to their differentiation into cholangiocytes and hepatocytes, which encompasses the downregulation of expression of crucial niche factors and temporally coincides with the exit of hematopoietic cells towards BM microenvironment ^58^. The reduction in abundance and downsizing of the spatial coverage of niche components occurs homogeneously throughout the FL parenchyma. Thus, at late stages of FL hematopoiesis, HSCs and HPCs become significantly less frequent but do not retract to specific sites and remain randomly distributed within FL.

While ECs and MSCs are relevant sources of HSC and HPC specific molecular cues, our findings point to a preeminent role of hepatoblasts in modulating early FL hematopoiesis. Among stromal cells, hepatoblasts express the highest levels, of *Kitl*, *Igf2* and *Angptl3*, as well as pro-differentiation cytokines such as *Epo* and *Thpo*, whose expression is shut down with differentiation. Our observations reinforce several previous studies, which noted the prolific cytokine secretion profile of hepatoblasts and their ability to maintain HSCs or HPCs in cocultures ^20–22,59^. Strikingly, a recent study reported no effects in FL and adult HSCs when *Kitl* was deleted from the hepatocyte lineage using *Alb*-Cre mice, thereby suggesting a minor contribution of hepatoblasts to the establishment of definitive hematopoiesis in FL ^60^. Nonetheless, to what extent Kitl is necessary for expansion of HSCs in the FL is yet unclear. Constitutive *Kitl* deficiency causes perinatal death due to anemia, which is most likely caused by impaired expansion of erythro-myeloid progenitors but not HSCs in the aorta gonad mesonephros and the FL ^35,61,62^. In turn, simultaneous targeting of *Kitl* in MSCs and ECs during embryonic development results in HSC deficiency after birth but not in the FL ^29^. Finally, our unpublished observations suggest inefficient targeting of hepatoblasts in *Alb*-Cre embryos at E13.5, when *Kitl* is most highly expressed. Therefore, further studies using refined *in vivo* strategies to selectively target hepatoblasts are needed to define the molecular basis of the potential regulatory role of this cell type on HSCs.

Beyond hepatoblasts, we find that ECs and MSCs express a wealth of crucial cytokines, which have been reported to not only regulate HSCs but also the maturation and differentiation of recently immigrated fetal HPCs. These findings are in line with the reported capacity of both cell types to support HSC expansion in vitro ^26,28,63^. Thus, FL HSC niches are multicellular and highly dynamic structures. This promiscuous and abundant expression of key factors in the stromal compartment points to a redundancy in niche support at molecular and cellular levels, which would ensure a robust function of FL throughout hematopoietic development and may explain why ablation of single cytokines from individual cellular sources does not cause strong suppressive effects on HSCs. Of note, the stromal compartment is likely more diverse and heterogeneous than resolved by our analysis. For instance, while stellate cells make up the largest fraction of the MSC fraction, other subpopulations have been described within the mesenchymal compartment of the FL. NG2^+^ periportal cells, which were also found to express factors such as *Kitl* and *Cxcl12*, are marked by Dlk-1 and CD140b, and thus are presumably included in the pool of cells here categorized as MSCs ^27^. These cells are nonetheless extremely rare at the onset of FL hematopoiesis, and spatially restricted to the adventitial layers of large portal vessels, when the full FL is pervaded with cells with very prominent HSPC supportive properties. In addition, mesothelial cells lining the external capsule of the FL, fall under the phenotypic signature of MSCs and are reported by an accompanying study as relevant sources for regulatory factors ^64^. Altogether, a more granular analysis of stromal elements is needed to unveil higher specialization of supportive functions at different stages of fetal hematopoiesis.

What drives the intense maturation and reorganization of the stromal infrastructure that underpins the exit of HSCs from the FL and the shift towards BM hematopoiesis? Our analyses uncover fundamental rearrangements in the transcriptional state of all stromal cells between E13.5 and E17.5, which reflect the high level of plasticity of the tissue infrastructure in this developmental period. In hepatoblasts, changes are largely linked to pathways previously described to mediate expansion and differentiation of this cell type, such as Wnt and Osm ^65–67^. Yet, the most striking observation is the notorious activation of inflammatory pathways found across all three cell types under sterile conditions in this short time window. Our computational analyses predict a number of likely molecular triggers of these emergent signatures in HSCs and stroma, which include IFN and TNF, as well as other inflammatory mediators such as IL-11, IL-4 and IL-27. Pro-inflammatory signaling has been shown to crucially influence HSC ontogeny, from emergence of definitive HSCs in the AGM to the specification of adult HSC programs ^68^. For instance, a recent study detected a spike of type I IFN signaling exactly during the timeframe investigated here, which contributes to the acquisition of mature gene expression programs in HSCs ^46^. Beyond IFN, IL-1b signaling has also been reported to control HSC expansion in mid-gestation ^69^. Therefore, our results reinforce the notion that oscillations in basal sterile inflammation levels influence the development of HSCs, not only via direct signaling but also through the reshaping of the function and structure of stromal cells in the different hematopoietic niches.

In summary, our findings suggest that FL HSC niches are transient structures formed by complex consortia of stromal cells, which provide a non-limiting supply of resources to support the differentiation and expansion of HSCs and HPCs at the peak stages of FL hematopoiesis. This propitious microenvironment is rapidly dismantled leading to the shift of HSCs towards more favorable territories. The elucidation of the dynamic arrangements and the precise signaling layers activated in these niches could have major implications in the field of regenerative medicine.

## Acknowledgements

The authors would like to thank Angelina Oestmann for technical assistance with mouse colony management. P.M.H. received support from the Forschungskredit fellowship of the University of Zurich. This work was supported by grants from the Swiss National Science Foundation (310030B_166673/1) to M.G.M, and the Novartis Foundation for Biomedical Research, the Swiss National Science Foundation (31003A_159597/1) and Consolidator Grant from the European Research Council (ERC-Co2019 865803) to C.N-A.

## Author contributions

P.M.H. and A.V designed and performed experiments, analyzed all data, and wrote the manuscript. A.G. implemented customized analysis tools for image segmentation and spatial statistics. K.L. and S.I. performed experiments. K.A.Z performed computational transcriptomic analyses. T.Z and I.R performed and discussed spatial data analyses. T.Y. and T.N. contributed transgenic mouse models and scientific discussions. M.G.M. contributed with scientific discussions. C.N.-A. conceived and directed the study, analyzed data and wrote the manuscript.

## Data availability

All raw and processed data will be made available upon reasonable request to the corresponding author.

## Supplementary Figure Legends

**Supplementary Figure 1**. graphs depicting gene ontology (GO) terms enriched in DEGs between hepatoblasts (HEP) mesenchymal stromal cells (MSC) and endothelial cells (EC). Number of genes per GO term and p values are shown.

**Supplementary Figure 2**. (**A**) Expression of selected hematopoietic cytokines and growth factors in the stromal EC, HEP, MSC subsets and hematopoietic CD45^+^ subset at embryonic timepoints E13.5, E15.5, E17.5 and postnatal P1.5. Data are displayed as mean + SEM. (**B**) Concentration of KITL in FL at different embryonic developmental timepoints as measured by ELISA (n = 8 embryos from at least two different experiments). Statistical significance was analyzed using the Kruskal-Wallis test with Dunn’s posttest. *, P < 0.05; **, P < 0.01; ***, P < 0.001.

**Supplementary Figure 3**. (**A**) FC plots of dissociated fetal livers at E13.5 of *Ctnnal1*^GFP/wt^ *Hlf*^tdTom/wt^ versus wild type controls. (**B**) Immunostained FL from a *Hlf*^tdTom/wt^ E13.5 embryo. CD117 (c-kit) is shown in green, nuclear staining with DAPI in blue and tdTom staining in red. All TdTom^hi^ cells, identified by microscopy, co-express c-kit (CD117). (**C**) FC analysis of stromal *Ctnnal1*^GFP^ expression in the stromal, non-hematopoietic compartment of the FL. A rare subset of GFP^+^ cells is detected on the CD45^-^Ter119^-^ fraction of FL. At E13.5 CD31^+^ and CD140b^+^ cells express GFP. Lower panels: histograms displaying GFP intensity of stromal CD31^+^ endothelial cells or CD140b^+^ mesenchymal cells. (**D**) Optical section of an immunostained fetal liver section confirms *Ctnnal1*^GFP^ expression on some ECs lining large portal vessels and on a subpopulation of fibroblastic pericytes. Vasculature labelled by Lyve1 (white) and GFP in green. (E) Immunostained thick FL slices of Ctnnal1^GFP/+^ transgenic mice, highlighting differential GFP expression on portal vessels and a central vein. All vessels are labeled by Lyve1 (magenta) and arterial structures by α-SMA (yellow).

**Supplementary Figure 4**. (**A**) Representative image of a DAPI stained FL slice (left) and segmented DAPI+ surface, which defines the entire tissue volume. This segmentation strategy was employed define tissue boundaries in quantitative and spatial analyses in the manuscript. (**B**) Representative image of segmented vessels use a deep learning model trained with ground truth data. Image shows segmentation of Lyve1+ vessel outlines. Details of the performance metrics of the vessel segmentation models can be found in ^53,70^. (**C**) Examples of annotated HPCs and HSCs in one 3D representative image of a FL from a *Ctnnal1*^GFP/wt^ *Hlf*^tdTom/wt^ E13.5 embryo

**Supplementary Figure 5**. (**A-C**) Right panels: median fluorescence intensities (MFI) in the GFP (A), tdTom (B) and DAPI (C) channels quantified within ± 2µm of the spot center for 2180 HPCs and 179 HSCs selected from all 8 slices analyzed at two different developmental timepoints. Data are plotted using the Tukey method. Left panels: the median fluorescence intensity for all cells annotated per slice is shown. Dots connected by lines represent the MFI in tdTom and GFP channels calculated for cells in the same tissue slice. (**D**) Empty space distance transform towards annotated progenitor cells shows widespread distribution of progenitor cells at a tissue scale level. Statistical significance was analyzed by Mann-Whitney *U* test with Dunn’s posttest for left panels in (A-C) and the Wilcoxon test for right panels *, P < 0.05; ** P < 0.01 ****, P < 0.001

## Methods

### Animal studies

Animals were maintained under standard conditions at the animal facility of the University Hospital Zürich and treated following the guidelines of the Swiss Federal Veterinary Office. All animal studies have been approved by the veterinary office of the Canton of Zürich. C57BL/6JRj mice were purchased from Janvier Labs (France), Kitl^GFP^ (Kitl^tm1.1Sjm^) and α-catulin^GFP^ (Ctnnal1^tm1.1Sjm^) mice were imported from The Jackson Laboratory (USA), Cxcl12^GFP^ mice (Cxcl12^tm2Tng^) provided by Takashi Nagasawa and Hlf^tdTom^ mice provided by Tomomasa Yokomizo. All transgenic mice have been previously _described_ ^32^,51,52,71.

### Tissue extraction, sectioning and immunostaining

Female mice (6-12 weeks) were raised in groups of five and 72 hours before mating exposed to bedding material from a male cage. In the evening (after 5 p.m.) one female each was transferred to a cage containing a single housed male. The consecutive morning (before 9 a.m.) the breeding pair was separated, and females analyzed for the presence of a vaginal plug. That timepoint was designated as E0.5. Females were only mated for one night and 12 days later examined for a possible pregnancy. Embryos were extracted from the previously euthanized pregnant dams through a small abdominal incision and further dissected under a stereo microscope (Leica). Fetal livers were dissected using forceps and separated into single lobes using a surgical scalpel under a stereo microscope the lobes were fixed in 2 % paraformaldehyde (diluted in PBS) (6 h, 4 °C), washed twice in ice-cold PBS for five minutes and subsequently dehydrated in 30 % sucrose in PBS (24 – 48 h, 4 °C). The liver lobes were placed into base molds (15×15×5 mm) and completely covered in OCT medium, snap-frozen using liquid nitrogen and stored at –80 °C until use.

The frozen OCT blocks were placed into 5 mL PBS for 15 minutes at room temperature until all OCT dissolved and successively washed twice in 5 mL PBS for 15 minutes to rehydrate the tissue. The liver lobes were again placed into base molds, excess liquid removed using a pipette and covered in 5 % low gelling temperature agarose (ca. 35 °C) and kept at 4 °C overnight. The solidified agarose block was cut into 240 µm thick transversal slices using a vibratome (Leica). Only centralized slices of each lobe were collected and moved into a 24-well plate using a soft paint brush. The slices were incubated in 400 µL blocking solution (0.2 % Triton X-100, 10 % donkey serum, in PBS) overnight, subsequently transferred into a new well containing 400 µL primary antibodies diluted in blocking buffer for 2 days, washed three times for 1 hour in wash solution (0.2 % Triton X-100 in PBS), incubated with secondary antibodies diluted in blocking solution for 2 days and successively washed three times for 1 hour in wash solution. All immunostaining steps were performed at 4 °C.

### Sample preparation, image acquisition and post processing

The liver slices are optically cleared by complete immersion into RapiClear for 24 hours, transferred onto a glass slide encompassing an iSpacer (250 µm), excess RapiClear removed by pipetting and sealed with a coverslip. Imaging was performed on a Leica SP8 confocal microscope system equipped with a white light laser source. Large tile scans were stitched using the Leica Application Suite (LAS) X software (Leica). Imaris software (v9.50, Oxford Instruments) was used for 3D visualization, post-processing and cell labeling. A median filter (3×3×3 pixels) was applied to channels containing HSC and progenitor labeling in order to reduce pixelate noise and simplify manual cell annotation.

### Volumetric quantitative image analysis

HSC and progenitor cells were manually annotated using the spots module of Imaris software (v8.40, Oxford Instruments) to ensure a minimal error rate. The segmentation of vasculature was achieved using an approach employing convolutional neural networks (modified U-Net), which were extensively trained with and evaluated on manual annotation data ^53^. Manual annotation and corrections of minor erroneous segmentation was performed using microscopy image browser (MIB Matlab 2.601) ^72^. Spatial statistics and analysis of cellular interactions has been previously described in ^42^. Briefly, the ESD (empty space distance) was calculated in order to describe all spatial constraints caused by the different segmented structures and cellular components. The ESD assigns every voxel within the analyzed tissue volume the shortest distance to the most proximal segmented structure. Voxels within intravascular regions were excluded from the analysis. Represented as a CDF (cumulative distribution function), the ESD describes the empirically measured fraction of tissue volume that is contained within any given distance to a segmented structure. For the analysis of the nearest neighbor distances, the distance to the nearest cell of the same type (HSC or HPC) was calculated based on the centers of the cells. To simulate a null distribution without any spatial clustering, the same number of cells was randomly placed within the segmented tissue, thereby sparing both types of vessels. Each simulation was repeated 500 times. Simulations and calculations were done using Mathematica 13.1 (Wolfram Research Inc.).

### Stromal cell isolation

Dissected FLs were placed into 5 mL digestion medium (DMEM GlutaMAX, 10 mM HEPES, 10 % fetal bovine serum (FBS) in a 6-well cell culture dish, disintegrated by gentle pulling motions using two micro-forceps, collagenase (Type 2, 0.04 g/mL) and Deoxyribonuclease I (0.2 mg/mL) added and thoroughly resuspended using a 1 mL pipette. The cell suspension was immediately incubated for 45 minutes at 37 °C with gentle sample agitation. Following incubation, 5 mL of ice-cold calcium– and magnesium-free phosphate-buffered saline (PBS) containing 10 % FBS were added, the suspension filtered through a 70 µm cell strainer, centrifuged at 500 g (5 minutes, 4 °C), the supernatant discarded, and the pellet resuspended into a single cell solution in PBS.

### Flow cytometry analysis

Single cell suspensions were incubated with TruStain fcX^TM^ (5 minutes, 4 °C) and subsequently immunostained using pre-conjugated antibodies (30 minutes, 4° C). All employed antibodies are listed in Table 1, including commercial sources. Next, the cells were washed twice in ice-cold PBS and resuspended in PBS containing DAPI (0.5 mg/mL), analyzed on a LSR II Fortessa (BD Biosciences). Post-acquisition data analysis was performed using FlowJo 10 software.

### Cell sorting, RNA isolation and RNA sequencing

Immunostained single cell solutions were prepared following the flow cytometry protocol explained above. Phenotypically defined stromal cell populations were separated using a FACS Aria (BD Biosciences) and collected directly into RNase-free microfuge tubes containing 400 µL RLT lysis buffer (QIAGEN) supplemented with 4 µL β-mercaptoethanol, and frozen down on dry ice for storage at –80 °C. Between 6’000 and maximal 50’000 cells were collected depending on the abundance of each cell type. RNA was extracted from thawed cell lysates using the RNeasy Plus Micro Kit (QIAGEN) following the manufacturer’s instructions and genomic DNA depleted using the supplied gDNA eliminator columns. RNA quality was judged by RIN values measured on a Bioanalyzer 2100 (Agilent, Waldbronn, Germany). Every analyzed sample displayed a RIN value above 9. Library preparation, cluster generation, and sequencing were performed at the functional genomics center Zurich (FGCZ, Zurich, Switzerland). The libraries were prepared using the SMART-Seq2 protocol (Picelli et al., 2014). Sequencing was performed on the Illumina HiSeq 4000 (single end 125 bp) using the TruSeq SBS Kit v4-HS (Illumina, Inc, California, USA).

### Bioinformatic analyses

Quality control and processing of RNA sequencing data was achieved using the SUSHI platform provided by the FGCZ, Zurich ^73^. The following integrated apps and settings were used: FastQC, FastQ Screen, STAR alignment to GRCm38.p5, featureCounts (Liao et al., 2014) and EdgeR for differential expression analysis adding a background expression of 10 reads.

Bulk RNA Sequencing DE analysis and heat map generation: Differentially expressed genes (DEGs) were calculated on the normalized count tables between conditions of interest and defined based on logFC cutoff > 1 or < 1 and adjusted p value < 0.05. For the heat map of marker genes for DLK, PDG and END scaled logCPM values were used. To identify those genes each cell type was compared against the other two and the overlapping genes were selected. Top selected marker genes were plotted on the heat map. For the box plot of selected genes at FL13, CPM values were used. Gene ontology analysis of differentially expressed genes with logFC > 1 was performed with clusterProfiler package and enrichGO function.

Pathway analysis: Pathway analysis was performed using two R packages PROGENy and decoupleR. PROGENy contains pathways and their target genes with weights for each interaction ^44^. For the analysis, we retrieved top 100 responsive mouse genes ranked by p value. To infer pathway activities Weighted mean method (wmean) was used. As input for wmean, DEGs for each cell type with absolute value logFC > 1 and adjusted p value below 0.05 were filtered. Log2FC of these DEGs was used to infer pathway activities between the two time points E13.5 and E17.5 in the HEP, MSC and EC. We also further visualized the most responsive genes with their logFC in the top activated pathway for each cell type.

NicheNet Ligand-Receptor Analysis ^45^: We first defined potentially active ligands, affected geneset of interest (DEGs as defined before according to log2FC and adjusted p value) and non-affected genes (background genes). NicheNet ranked ligands according to how well they predict whether a gene belongs to the gene set of interest compared to the background gene set. As potentially active ligands we considered all ligands in the NicheNet model that were found in our data set after removing lowly expressed genes. Next, we ordered the potential ligands according to their activity (by Pearson correlation) in all cell types and selected top 6 ligands. Visualization of the activity of these ligands was done in a heatmap depicting z-scores. Then, we determined the receptors for the selected ligands using the receptors in the NicheNet model that were expressed after filtering for lowly expressed genes. These receptors are shown in the heat map with their z-scores. We also show the weight of the interaction between the ligand and receptor in the integrated weighted ligand signaling network of NicheNet. For the heat map of target genes regulated by the ligands, we selected genes with the highest regulatory potential and highest logFC between time points within each cell type.

### Bioinformatic analyses

Statistical analyses were performed using Graphpad Prism. The statistical tests employed are indicated in the figure legends. Graphs depict mean ± SD unless stated otherwise.

### Supplemental Movie Legends

**Movie S1. Multiscale volumetric imaging of FL lobes.** Volumetric representations of a thick slice of an immunostained fetal liver lobe of a E13.*5 Cxcl12*^GFP/wt^ embryo at multiple scales and resolutions. Tissue-wide low magnification image (left) shows widespread distribution of GFP^+^ (green) cells throughout the FL. Sinusoidal vasculature marked by Lyve-1 expression (magenta) and large portal vessels are surrounded by a layer of αSMA^+^ cells (yellow). Higher magnification images and dynamic rastering throughout the z-axis illustrates the visualization capabilities of this technique and the detailed morphological features of GFP-expressing cells.

**Movie S2. Timeline analysis of hepatic Cxcl12 expression in Cxcl12^GFP/wt^ embryos, pups and adult mice.** Volumetric representations of thick feta liver slices covering entire lobes are shown. Early embryonic stages are characterized by intense stromal GFP expression and only sparse GFP expressing cells can be detected in adult livers in proximity to large vasculature. Embryonic day (E), postnatal day (P). Lyve-1(magenta) αSMA (yellow), GFP (green).

**Movie S3. tdTomato allows visualization of c-kit^+^Hlf*-tdTom*^+^ HPCs in FL from slices.** Representative example of a FL slice from a *Hlf*^tdTom/wt^ E13.5 embryo stained for tdTom (red), c-kit (green) and Lyve-1 (white). Image shows the high density of c-kit-expressing progenitors in the FL at these stages. A limited fraction of c-kit^+^ cells is additionally marked by *Hlf-*tdTom expression. Double positive cells constitute earliest multipotent progenitor cells.

**Movie S4. Visualization of HSCs and HPCs in the FL *Ctnnal*1^GFP^*Hlf*^tdTom^ double reporter mice.** Representative tissue-wide image of a FL from a E13.5 *Ctnnal*1^GFP^*Hlf*^tdTom^ embryo stained for GFP (green), tdTom (red), DAPI (blue) and Lyve-1 (magenta). Examples of HSCs expressing both reporters depicted in yellow (mixed expression of GFP and tdTom) and HPCs (GFP^-t^dTom^+^) are shown scattered throughout the entire FL lobe.

