## Supplementary figures and images for "Tissue-scale dynamic mapping of Hematopoietic Stem Cells and supportive niche cells in the fetal liver"

### Supplemental Figure 1

Supplemmentary Figure 1

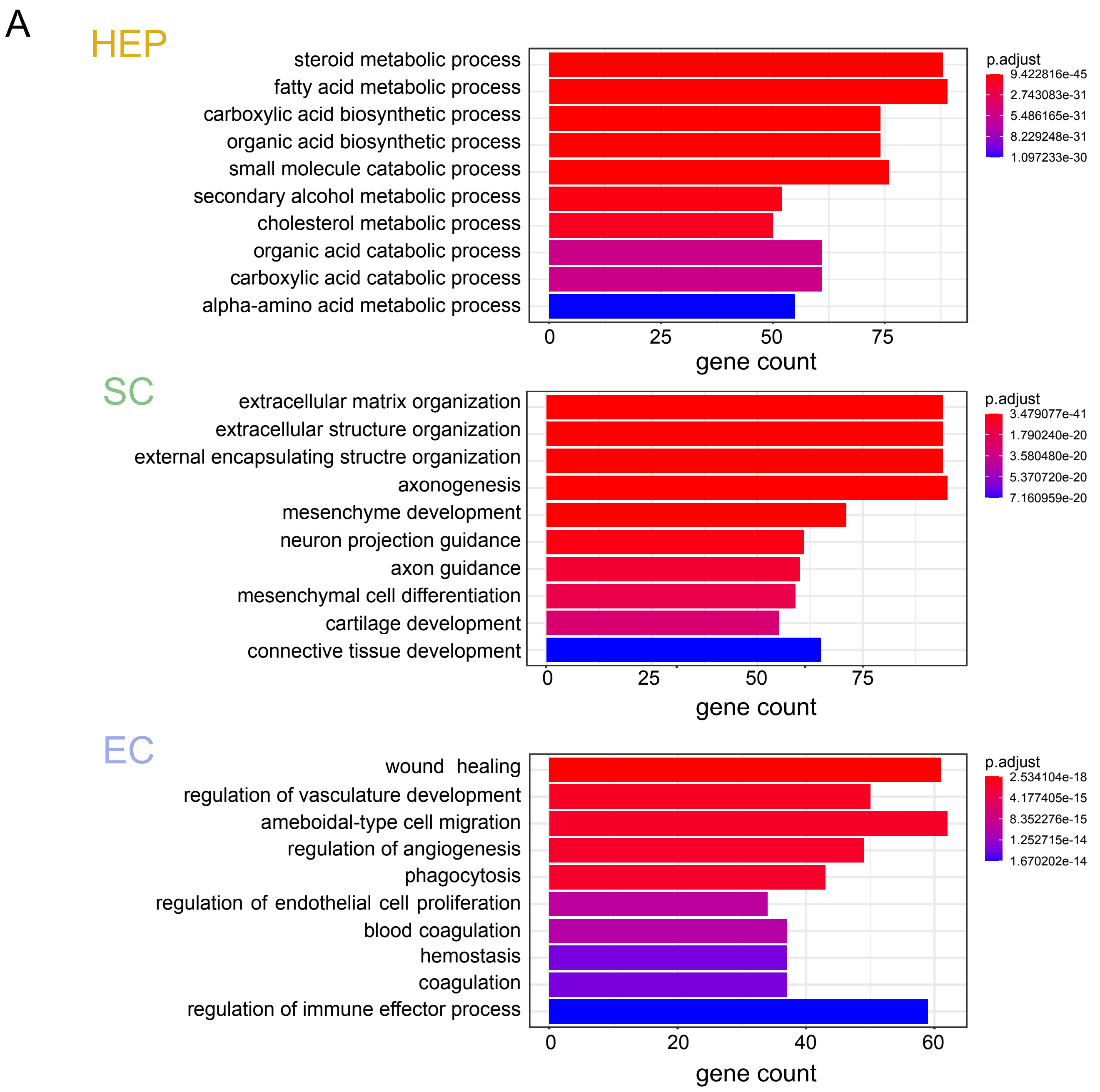

### Supplemental Figure 2

#### Supplemmentary Figure 2

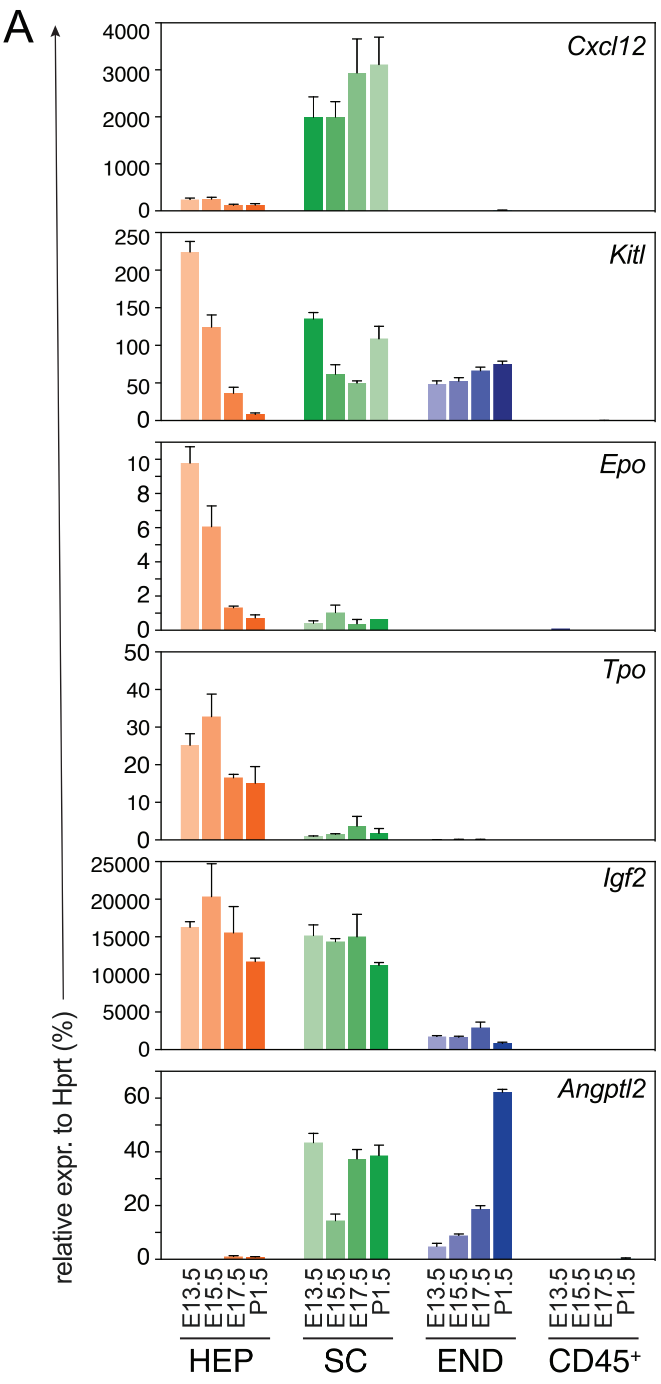


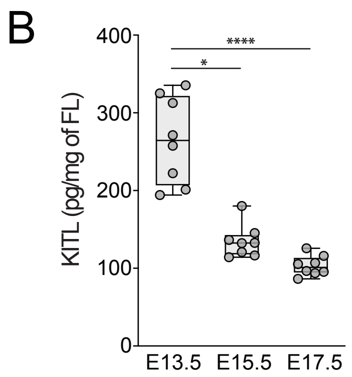

### Supplemental Figure 3

#### Supplementary Figure 3

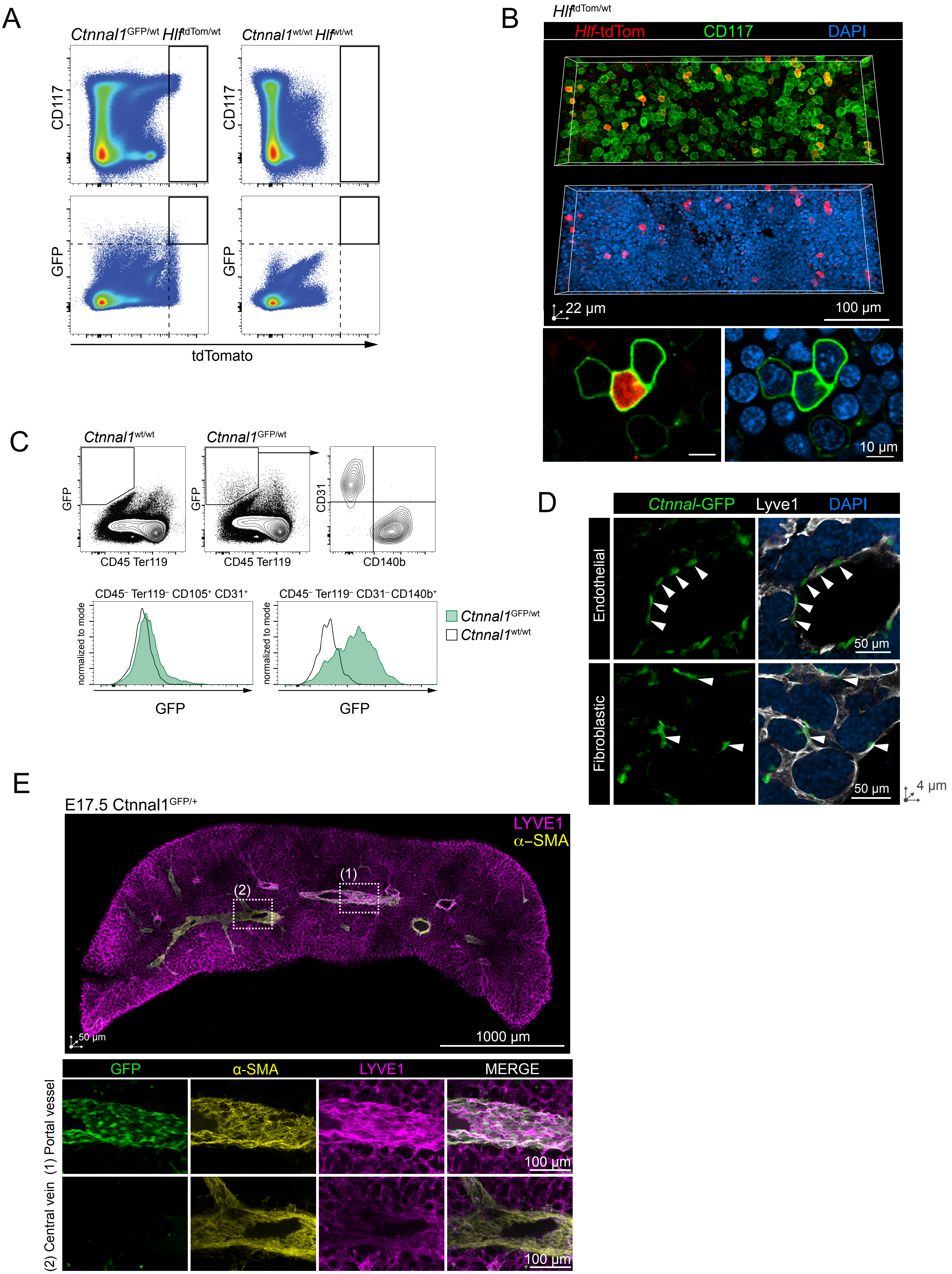

### Supplemental Figure 4

#### Supplemmentary Figure 4

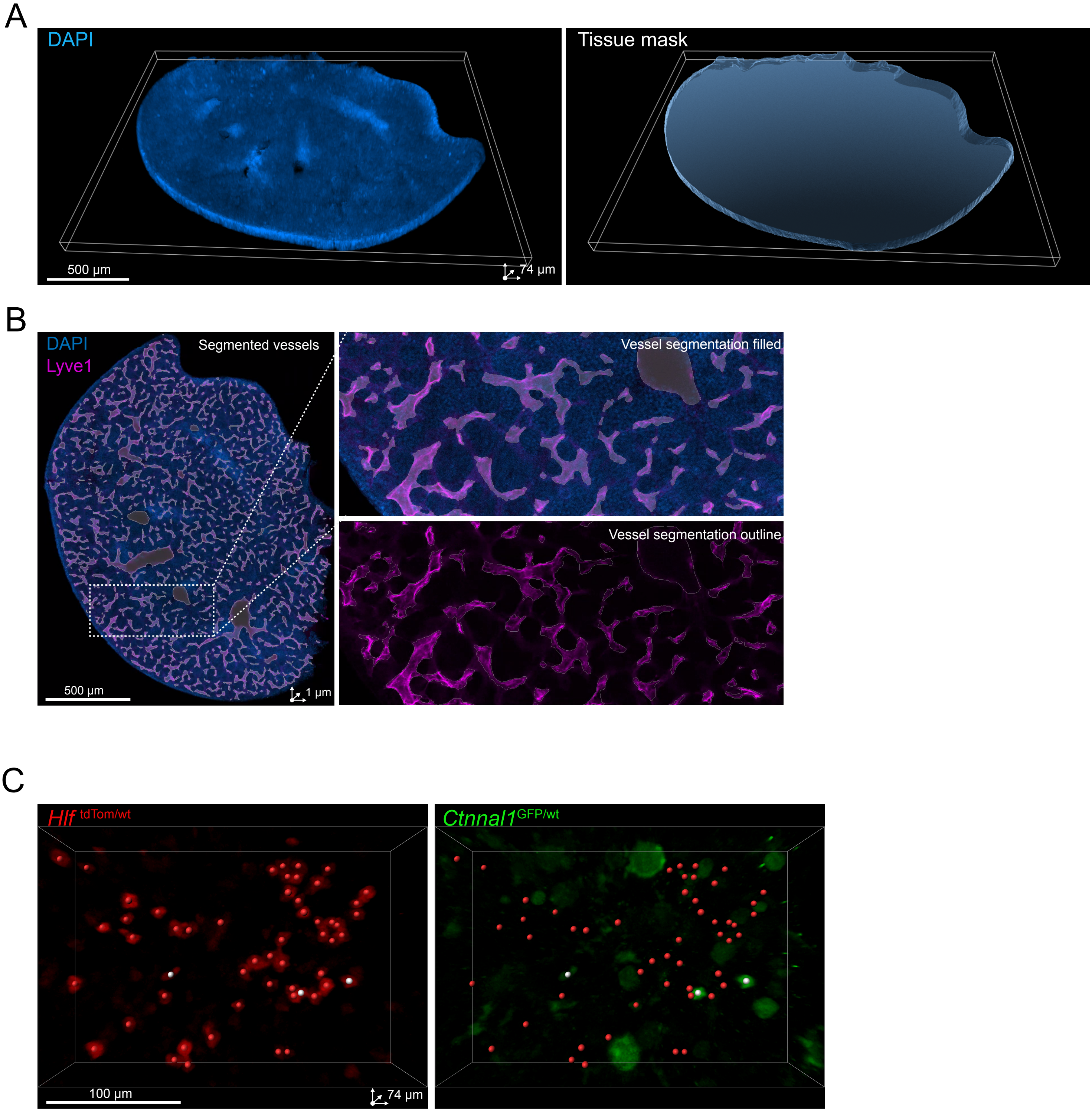


Annotated spots Hlf⁺ Ctnnal1⁻ (HPCs)
Annotated spots Hlf⁺ Ctnnal1⁺ (HSCs)

### Supplemental Figure 5

#### Supplemmentary Figure 5

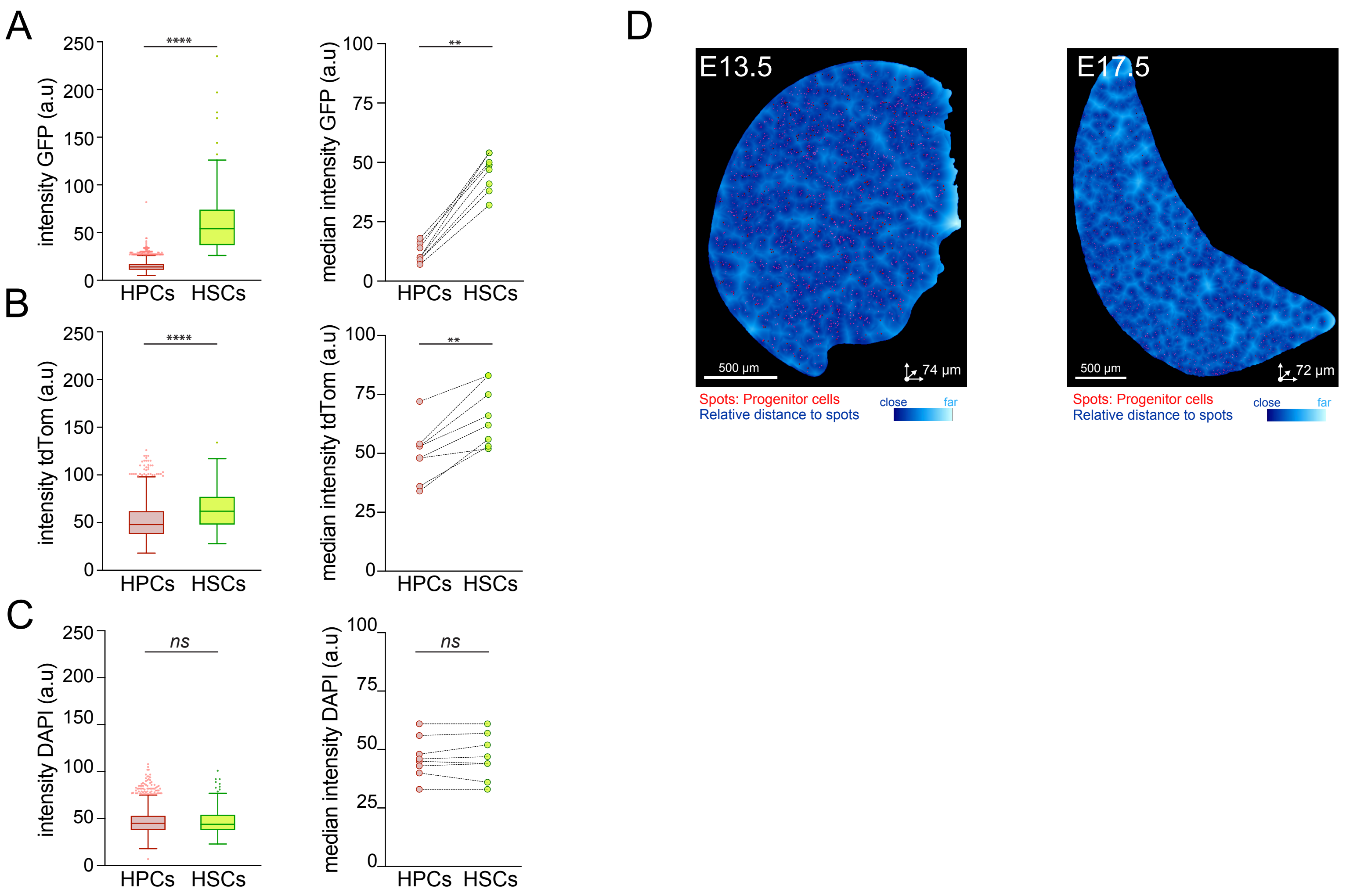
